# Trailer Hitch tunes condensate phase behavior and mRNA partitioning to regulate P-body homeostasis during *Drosophila melanogaster* oogenesis

**DOI:** 10.1101/2025.06.18.660414

**Authors:** Samantha N. Milano, Livia V. Bayer, Julie J. Ko, Gwendolyn S. Posner, Caroline E. Casella, Diana P. Bratu

## Abstract

Processing bodies (P-bodies) are cytoplasmic granules that regulate mRNA storage, repression, and decay, yet how their internal organization supports selective mRNA regulation remains poorly understood. Here, we show that the conserved LSm protein Trailer Hitch (Tral) is a key organizer of P-body architecture and function in the *Drosophila melanogaster* female germline. Using quantitative confocal imaging, super-resolution microscopy, and chemical perturbation of intermolecular interactions, we demonstrate that Tral coordinates the incorporation and spatial organization of the core P-body proteins Me31B and Cup. Loss of Tral alters their partitioning into P-bodies, promotes demixing into distinct subdomains, and shifts condensates toward a less dynamic, structurally heterogeneous state. These organizational changes have functional consequences for mRNA storage: Tral depletion selectively releases maternal mRNA *bicoid*, while *nanos* mRNA remains P-body associated and stable. We further identify *twinstar* mRNA, encoding the actin regulator Cofilin, as a Tral-dependent P-body client whose localization and organization within P-bodies requires Tral:RNA interactions and electrostatic forces. Reduced *twinstar* mRNA levels in the absence of Tral are associated with decreased nuclear G-actin and altered transcription of *me31B* and *cup*, revealing a potential feedback mechanism that links cytoplasmic P-body organization to nuclear gene expression. Together, these findings establish Tral as a central regulator of P-body architecture that couples condensate organization to selective mRNA regulation and transcriptional homeostasis.

## INTRODUCTION

Membraneless organelles function in a multitude of mRNA processing pathways ranging from transcript storage to degradation^1–3^. These granules are defined as liquid-like bodies which are separated from the surrounding cytoplasm via liquid-liquid phase separation (LLPS)^4^. LLPS occurs when specific proteins and mRNAs become locally concentrated in the cytoplasm and reach a threshold at which it becomes thermodynamically favorable for a distinct liquid phase to emerge. This process can be spurred by post-transcriptionally regulated mRNAs which act as scaffolds for RNA-binding proteins (RBP)^5^. Many RBP have intrinsically disordered regions (IDRs) which can also initiate phase separation as well as tune the emergent properties of condensates^6–8^. Once LLPS granules nucleate, they can grow by fusion with other liquid-like granules via a process called ‘Ostwald ripening’ and they can remain liquid-like by continually breaking down intra-condensate bonds^9,10^. Over time, these granules can mature into gel-like condensates which are defined by slower fusion rates and slower internal recovery after FRAP (fluorescence recovery after photobleaching) analysis. These granules are less dynamic, but their arrested state can provide additional nuance to condensate function^11^.

Processing bodies (P-bodies) are one such tunable LLPS granules. P-bodies are defined by their constitutive presence in the cytoplasm as well as their active involvement in mRNA processing^2,3,12^. P-bodies have been directly implicated in mRNA decay, with studies quantifying the process and demonstrating that decay efficiency is significantly enhanced in the presence of these granules despite their presence not being necessary for decay to occur^13,14^. Paradoxically, P-bodies also play a critical role in mRNA repression and storage, as the knockdown of key P-body proteins leads to ectopic expression of repressed transcripts^15,16^. Given the divergent functions of these granules, it has been difficult to elucidate how a single organelle can coordinate mRNA processing in a transcript-specific manner. Recently, a landmark study has highlighted how the condensate state of LLPS bodies contributes to mRNA association with granules, suggesting that emergent condensate properties can drive function^11^. This provides a mechanism by which P-body organization can facilitate specific mRNA regulation, but it remains unclear how P-bodies can autoregulate their organization and phase state in order to govern mRNA fate.

With the aim of unraveling how P-bodies self-organize to function in differential transcript storage, we chose to focus our study on the contributions of Trailer Hitch (Tral). Tral is a conserved LSm protein found in P-bodies with homologs across species^17^. It directly binds to mRNA making it an RBP, it has long IDRs, and is capable of direct protein binding^15,18,19^. In *D. melanogaster*, Tral is part of a central translational repression mRNP together with Me31B, an RNA helicase, and Cup, an eIF4E-binding protein. Tral and Me31B bind directly to repressed transcripts and work to prevent the recruitment of the mRNA decay machinery^15,20^. Cup is a large intrinsically disordered proteins that facilitates translational repression by out-competing the binding of the translation initiation complex ^21^. Tral lies at the center of this mRNP as it can bind directly to Me31B via an FDF domain as well as directly to Cup, via its LSm domain^18,22^.

Tral also appears to play a separate role in actin regulation. Tral knockdown egg chambers develop actin cages which collect secreted proteins in the oocyte^23,24^. Furthermore, Tral mutants display a *dumpless* phenotype, where nurse cell nuclei get stuck at the ring canals which is largely a result of mis-regulated actin dynamics^24^. Knockdown of Cup or Me31B does not result in these phenotypes indicating that Tral’s role in the actin life cycle may be independent of its P-body function.

Over the past two decades, the role of actin has expanded significantly, with mounting evidence now confirming that actin is present in the nucleus, and it plays diverse and essential roles in transcriptional regulation^25^. Nuclear actin is crucial for the assembly of the pre-initiation complex, contributes to chromatin remodeling, facilitates histone modifications at transcription sites, and modulates the binding efficiency of transcription factors, positioning it as a pivotal regulator of gene expression^26^.

In this study, we uncover a novel role for Tral in contributing to nuclear actin homeostasis by regulating *twinstar* mRNA, which in turn appears to affect the transcription rates of *me31B* and *cup* mRNAs, thus providing a mechanism for the interdependent regulation of core P-body protein levels via a cytoplasmic sensor. Using super-resolution imaging and RNAi-mediated knockdowns, we also show that Tral is essential for the proper organization of Me31B and Cup within P-bodies and demonstrate that mRNA storage in P-bodies is regulated by the synergistic interactions between Tral, Cup, and Me31B, as disrupting these interactions, either by Tral knockdown or through chemical treatments, differentially affects transcript storage. Interestingly, while Tral knockdown leads to the release of *bicoid,* a maternal mRNA, the association of *nanos*, another maternal mRNA with P-bodies remains unaffected, and *twinstar* mRNA aberrantly organizes within P-bodies, highlighting that mRNAs are differentially regulated within P-bodies by diverse intramolecular interactions.

## RESULTS

### Tral differentially affects the incorporation of Me31B and Cup into P-bodies

The *D. melanogaster* female germline is the ideal tissue to study P-body self-regulation and function as it epitomizes the need for precise spatiotemporal localization of mRNA transcripts over long distances. *D. melanogaster* ovaries are made up of ∼18 ovarioles each of which is composed of developing egg chambers progressing from the germarium, where the germline stem cells are housed, into the oviduct from which they are eventually deposited. Each germline stem cell divides to create a daughter cell which then undergoes 4 rounds of mitosis with incomplete cytokinesis. This results in a 16-cell egg chamber which shares a common cytoplasm through the F-actin ring canals that are left behind by the incomplete cytokinesis. One of these 16 cells is designated as the oocyte and is predominantly transcriptionally silent throughout oogenesis^27^. As a result, all maternally deposited mRNAs required for development are transcribed in the other 15 cells (nurse cells) and must be stably maintained and repressed as they are transported through the ring canals and into the oocyte where they are precisely localized (Fig. 1A)^28,29^. While the function of P-bodies in the oocyte is complex, requiring the differential storage, localization, and release of diverse maternal mRNAs, P-body function in the nurse cells is comparatively straightforward, as all maternal mRNAs must be safely stored and transported to a single destination: the oocyte. This simplicity makes nurse cells an ideal system for studying the fundamental mechanisms by which P-bodies self-organize to facilitate mRNA storage.

**Figure 1:**
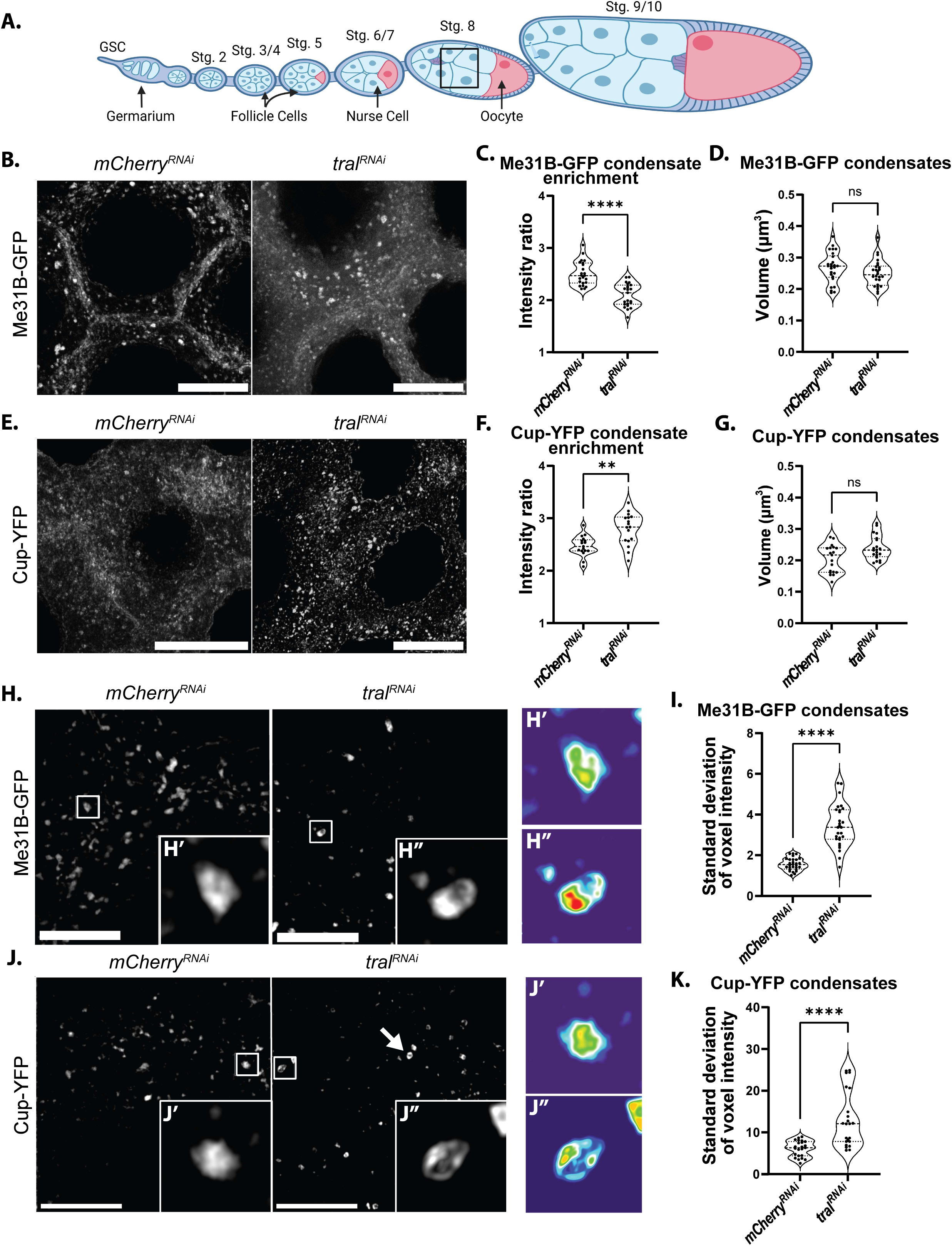
Tral differentially affects the incorporation of Me31B and Cup into P-bodies. **(A)** Schematic of a fly ovariole. Developmental stages progress from left to right. The oocyte (pink) and the nurse cells (light blue) are surrounded by somatic follicles (darker blue). The outlined ROI (black square) indicates the proximate region of the nurse cells visualized in all zoomed-in image panels. **(B)** Nurse cells expressing endogenous Me31B-GFP in *mCherry^RNAi^* (control) and *tral^RNAi^* egg chambers. Images are XY projections of 5 optical Z slices of 0.3µm. Scale bars are 20µm. **(C)** Intensity ratio measurements for **(B)**; average fluorescence intensity of condensates was divided by cytoplasmic fluorescence intensity to provide condensate enrichment in *mCherry^RNAi^* and *tral^RNAi^* egg chambers (n = 23). **(D)** Volume size quantifications comparing Me31B-GFP condensates in *mCherry^RNAi^* and *tral^RNAi^* egg chambers (n = 26). **(E)** Nurse cells expressing endogenous Cup-YFP in *mCherry^RNAi^* and *tral^RNAi^* egg chambers. Images are XY projections of 5 optical Z slices of 0.3µm. Scale bars are 20µm. **(F, G)** Same as **(C, D)** for Cup-YFP (n = 17 and n = 19, respectively). **(H)** Visualization of endogenous Me31B-GFP via STED. Images are XY projections of 5 optical Z slices of 0.22µm. Scale bar is 10µm (2µm for the zoomed inset). **(H’, H”)** Fluorescence intensity heat maps of a P-body in *mCherry^RNAi^* and *tral^RNAi^* egg chambers, respectively. **(I)** Standard deviation of voxel fluorescence intensity values for Me31B-GFP condensates in *mCherry^RNAi^* and *tral^RNAi^* egg chambers (n = 28). **(J-K)** Same as **(H-I)** for Cup-YFP. Arrow indicates Cup-YFP ring structure (n = 23). For all plots, each data point represents the average value of all P-bodies detected in an image. Significance was assessed using Mann-Whitney statistical tests. Error bars represent standard deviation. **** P < .0001.

The storage of many maternal mRNAs is regulated through the coordinated actions of Tral, Me31B, and Cup. Notably, these three proteins also contribute to the structural integrity of P-bodies across species, and consistent with this, in our control *mCherry^RNAi^* egg chambers, all three proteins were colocalized within P-bodies (Fig. S1A)^30–32^. Since Tral can directly interact with both Me31B and Cup, we wanted to examine whether loss of Tral alters their assembly into condensates, potentially affecting mRNA storage. To address this, we utilized the UAS/Gal4 system to drive expression of *tral^RNAi^* while covisualizing endogenously tagged Me31B-GFP and Tral-RFP with immunolabeled Cup (Fig. S1A). Tral knockdown led to a reduction of *tral* mRNA and Tral-RFP protein levels confirming RNAi efficiency. Despite changes in Me31B-GFP and Cup cytoplasmic distributions, they remained colocalized in the *tral^RNAi^* background (Fig. S1A,B).

To quantify these changes and their enrichment in condensates, we divided the average protein fluorescence intensity in condensates with the average intensity in the cytoplasm in nurse cells of mid-stage (7-8) egg chambers (Fig. 1A, black square). Interestingly, we found that in Tral knockdown egg chambers, ∼16% less of the available Me31B-GFP partitioned into P-bodies, while the average condensate volume remained unchanged (Fig. 1B-D). However, condensate fluorescence intensity decreased by ∼21%, collectively indicating an alteration in P-body structure (Fig. S1C). To verify the altered distribution of Me31B-GFP, we generated 3D cytoplasmic fluorescence intensity maps under both conditions and quantified signal heterogeneity by calculating the standard deviation of voxel intensities. A more punctate distribution corresponds to a higher standard deviation, whereas a more diffuse distribution results in a lower value. We observed a ∼47% decrease in standard deviation in *tral^RNAi^* egg chambers, consistent with a markedly more diffuse Me31B-GFP distribution (Fig. S1D). Next, we visualized endogenously tagged Cup-YFP and found the opposite trend with ∼13% more of the available Cup-YFP partitioned into P-bodies, while the volume of Cup-YFP condensates was again unaffected (Fig. 1E-G). Here too, the fluorescence intensity of the condensates increased by ∼76%, again suggesting an altered P-body internal organization (Fig. S1E). This interpretation was further validated by analysis of a 3D cytoplasmic fluorescence intensity map, in which the standard deviation of voxel intensities increased by ∼43%, indicating a more heterogeneous (punctate) cytoplasmic distribution (Fig. S1F).

To gain a deeper understanding of Tral’s effect on intracondensate structure, we employed STED super-resolution imaging and assessed the distribution of fluorescent proteins within individual condensates. By determining the fluorescence intensity of each voxel within a condensate and quantifying the standard deviation of these voxel intensities, we were able to quantify condensate organization. Condensates with highly organized, solid-like structures typically exhibit regions of high intensity (fluorescent peaks) and regions of low intensity (fluorescent valleys), resulting in a high standard deviation, which is characteristic of a rough surface. In contrast, more liquid-like condensates, where fluorescently tagged proteins are able to flow within the granules, display more uniform voxel intensities, leading to lower standard deviations, which is indicative of a smooth surface^33,34^. Using this approach, we found that Me31B-GFP condensates in the Tral knockdown background had ∼120% higher standard deviations of voxel intensity when compared to the *mCherry^RNAi^* control background, indicating that Me31B-GFP condensates are rougher in Tral knockdown backgrounds, suggesting a more solid-like phase state (Fig. 1H, I). Similarly, Cup condensates became ∼117% rougher, indicating more heterogeneous intracondensate structures and a more solid-like phase state (Fig. 1J, K). Notably, many of these condensates adopted a ring structure further hinting at altered internal organization (Fig. 1J, arrow). Collectively, these results indicate that Tral plays a role in orchestrating the incorporation and spatial arrangement of Cup and Me31B within P-bodies, potentially tuning their biophysical properties.

### Tral promotes a shared condensate state between Me31B and Cup

Given that Tral is essential for proper distribution of P-body components Cup and Me31B (Fig. 1H-K), we hypothesized that Tral facilitates key intramolecular interactions that support P-body organization and function. To test this, we used chemical treatments of live egg chambers to probe the molecular forces stabilizing the condensate states of individual P-body proteins. We quantified these effects by measuring condensate sphericity, as increased sphericity is a well-established hallmark of condensate breakdown resulting from the disruption of stabilizing interactions and leading to a more liquid-like granule^11,30^.

We first visualized the effects of 1,6-hexanediol, an aliphatic alcohol, known to disrupt hydrophobic interactions essential for phase-separated condensate integrity^35^. Control egg chambers expressing Me31B-GFP were treated with 1% 1,6-hexanediol for 30 minutes and subsequently fixed to visualize endogenous Me31B-GFP. This treatment led to increased sphericity of Me31B-GFP condensates in nurse cells, suggesting that under control conditions, their structural integrity depends on hydrophobic interactions, consistent with prior observations in late-stage oocytes (Fig. 2A, B)^11^. To test the contribution of electrostatic interactions, which also play a key role in P-body phase behavior, we treated *mCherry^RNAi^* egg chambers with 200mM NaCl^36,37^. This treatment similarly increased Me31B-GFP condensate sphericity, indicating a dual reliance on hydrophobic and electrostatic forces for condensate integrity (Fig. 2A, B).

**Figure 2:**
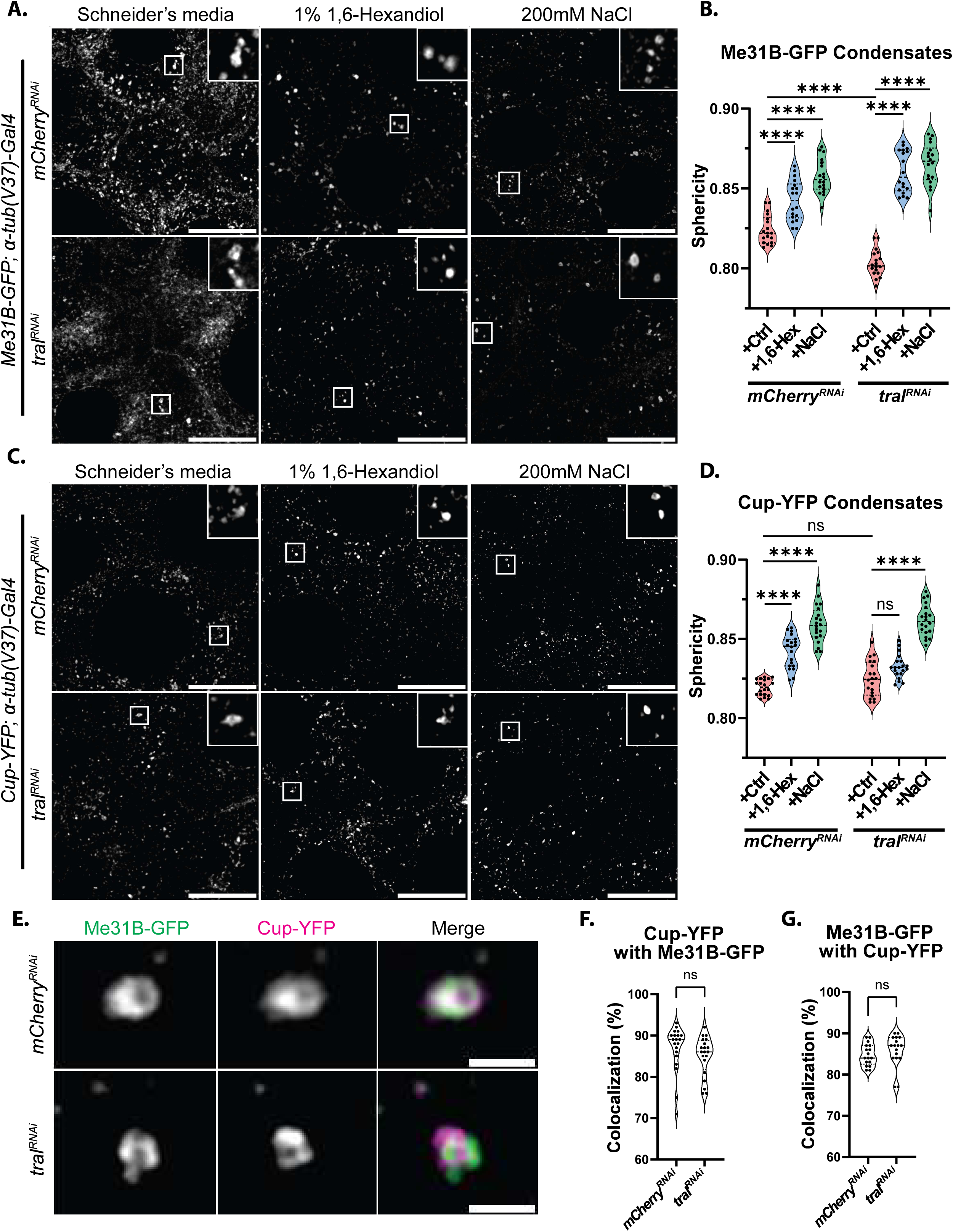
Tral promotes a shared condensate state between Me31B and Cup. **(A)** Me31B-GFP visualized in *mCherry^RNAi^* and *tral^RNAi^* egg chambers treated with Schneider’s media (+Ctrl), 1% 1,6-hexanediol, or 200mM NaCl. Images are XY projections of 5 optical Z slices of 0.3µm. Scale bars are 20µm. **(B)** Sphericity quantifications comparing Me31B-GFP labeled condensates in *mCherry^RNAi^* and *tral^RNAi^* egg chambers under conditions in **(A)** (n = 19). **(C)** Cup-YFP visualized in *mCherry^RNAi^* and *tral^RNAi^* egg chambers treated with Schneider’s media (+Ctrl), 1% 1,6-hexanediol, or 200mM NaCl. Images are XY projections of 5 optical Z slices of 0.3µm. Scale bars are 20µm. **D)** Sphericity quantifications comparing Cup-YFP labeled condensates in *mCherry^RNAi^* and *tral^RNAi^* egg chambers under conditions in **(C)** (n = 20). **(E)** STED images of Me31B-GFP with Cup-YFP. Images are XY projections of 5 optical Z slices of 0.22µm. Scale bars are 2µm. **(F, G)** Colocalization analysis of Cup-YFP with Me31B-GFP, and Me31B-GFP with Cup-YFP (n = 18 and n =17, respectively). For all plots, each data point represents the average value of all P-bodies detected in an image. Significance was assessed using Mann-Whitney statistical tests. Error bars represent standard deviation. **** P < .0001.

We next asked whether these dependencies were altered in the absence of Tral. Me31B-GFP condensates in *tral^RNAi^* egg chambers also became more spherical upon 1,6-hexanediol and NaCl treatment, indicating that both hydrophobic and electrostatic interactions continue to support Me31B phase behavior in the absence of Tral (Fig. 2A, B). Interestingly, we observed a significant reduction in sphericity in the control condition in the *tral^RNAi^* background compared to *mCherry^RNAi^*, suggesting that Tral is required to maintain Me31B condensate integrity, perhaps through its direct binding. This finding is consistent with our earlier observation that Me31B-GFP condensates appear rougher in the absence of Tral (Fig. 1H, I).

In *mCherry^RNAi^* egg chambers, Cup responded similarly to Me31B, becoming more spherical following both 1,6-hexanediol and NaCl treatment. Strikingly, however, Cup-YFP condensates in *tral^RNAi^* egg chambers similarly reacted to NaCl, but failed to respond to 1,6-hexanediol, suggesting that hydrophobic interactions were no longer involved in maintaining Cup’s phase state in the absence of Tral (Fig. 2C, D). Furthermore, we did not observe a statistically significant difference in sphericity between untreated *mCherry^RNAi^* and *tral^RNAi^* Cup-YFP condensates; however, there was a trend toward increased sphericity in the *tral^RNAi^* background, consistent with our earlier observation that Cup-YFP condensates appear rougher in the absence of Tral (Figs. 2D and 1J, K). As 1,6-hexanediol has possible off target effects, we repeated our analysis with SDS which similarly disrupts hydrophobic interactions^38^. Notably, we were able to replicate our previous findings with 0.5% SDS. Under control conditions, both Me31B-GFP and Cup-YFP condensates became more spherical and in the Tral knockdown background, SDS treatment led to Me31B-GFP marked condensates becoming more spherical, while Cup-YFP labeled condensates remained unaffected (Fig. S2A-D).

Together, these findings demonstrate that in wild-type egg chambers, both Me31B and Cup rely on hydrophobic and electrostatic interactions to maintain their condensate states. However, in the absence of Tral, Me31B remains sensitive to both forces, while Cup becomes exclusively reliant on electrostatic interactions. The loss of hydrophobic interactions, specifically for Cup, suggests that Tral plays a crucial role in coordinating the shared biophysical properties of Me31B and Cup, unifying their condensate behavior within P-bodies.

To further investigate this coordination, we employed STED super-resolution microscopy to visualize the intracondensate distribution of Me31B and Cup. In control egg chambers, Me31B-GFP and Cup-YFP largely overlapped within P-bodies. However, in *tral^RNAi^* egg chambers, the two proteins demixed and occupied distinct subdomains within the same P-body, indicating that Tral is required to maintain their unified phase state (Fig. 2E). Interestingly, although Cup and Me31B demixed within individual condensates, their overall colocalization at the level of P-body association remained unchanged (Fig. 2F, G). This suggests that Tral may organize their spatial distribution within the condensate, but not necessarily their recruitment.

### Tral is necessary for maintaining P-body function in select transcript storage

Given that Tral influences the condensate states of Me31B and Cup, we next examined whether the loss of Tral affects the storage of key maternal transcripts. We focused on two well-characterized maternal mRNAs, *bicoid* and *nanos*, which exemplify opposite ends of the maternal transcript expression spectrum: *bicoid* is stringently repressed throughout oogenesis, whereas *nanos* is expressed in the germarium and then again at late stages^39–41^.

Visualization of *bicoid* and *nanos* mRNAs in *tral^RNAi^* egg chambers revealed that *nanos* maintained normal colocalization with Me31B-GFP labeled P-bodies, whereas *bicoid* colocalization was reduced by ∼32% (Fig. 3A-C). A similar reduction in *bicoid* colocalization was observed with Cup-YFP marked P-bodies (Fig. S3A-C). To determine whether this loss of colocalization impacted transcript stability, we quantified total mRNA levels. Notably, *bicoid* mRNA abundance decreased by ∼18% in *tral^RNAi^*egg chambers, while *nanos* mRNA levels remained unchanged (Fig. 3D).

**Figure 3:**
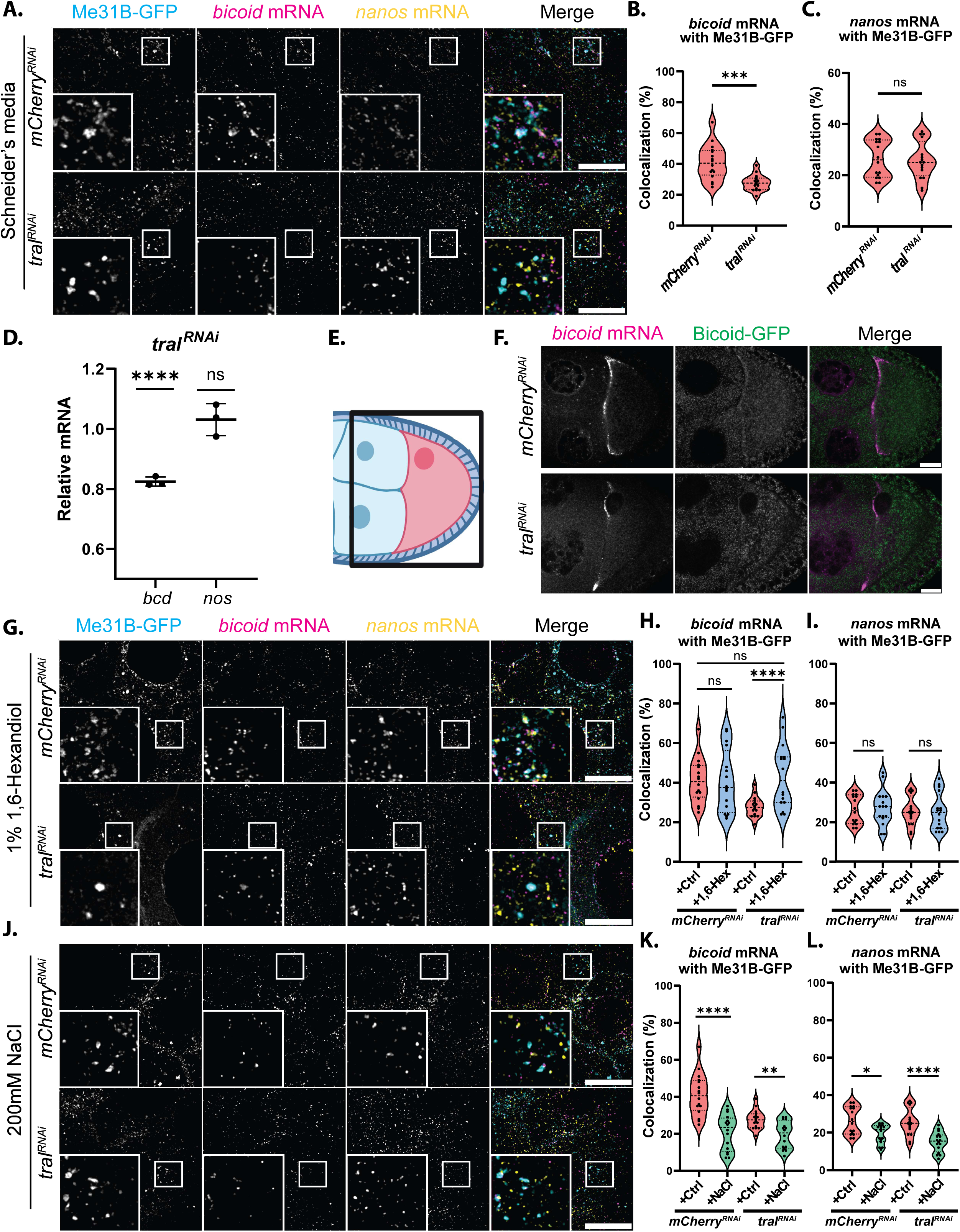
Tral is necessary for maintaining P-body function in select transcript storage. **(A)** Me31B-GFP covisualized with smFISH probes labeled *bicoid* and *nanos* mRNAs in *mCherry^RNAi^*and *tral^RNAi^* egg chambers incubated in Schneider’s media (+Ctrl). Images are XY projections of 5 optical Z slices of 0.3µm. Scale bars are 20µm. **(B)** Colocalization calculations for *bicoid* mRNA with Me31B-GFP condensates (n = 16). **(C)** Colocalization calculations for *nanos* mRNA with Me31B-GFP condensates (n = 16). **(D)** *bicoid* and *nanos* mRNA levels calculated with RT-qPCR in *tral^RNAi^* egg chambers. Significance calculated with Welch’s t-test (n = 3). **(E)** Diagram of ROI for egg chamber analysis in **(F).** **(F)** *bicoid* mRNA visualized with Bicoid-GFP in *mCherry^RNAi^* and *tral^RNAi^* egg chambers. **(G)** Me31B-GFP covisualized with smFISH probes labeled *bicoid* and *nanos* mRNAs in *mCherry^RNAi^* and *tral^RNAi^* egg chambers incubated in 1% 1,6-hexanediol. Images are XY projections of 5 optical Z slices of 0.3µm. Scale bars are 20µm. **(H)** Colocalization calculations for *bicoid* mRNA with Me31B-GFP condensates in (**G)** (n = 16). **(I)** Colocalization calculations for *nanos* mRNA with Me31B-GFP condensates in **(G)** (n = 16). **(J)** Me31B-GFP covisualized with smFISH probes labeled *bicoid* and *nanos* mRNA in *mCherry^RNAi^* and *tral^RNAi^* egg chambers incubated in 200mM NaCl. Images are XY projections of 5 optical Z slices of 0.3µm. Scale bars are 20µm. **(K)** Colocalization calculations for *bicoid* mRNA with Me31B-GFP condensates in (**J)** (n = 16) **(L)** Colocalization calculations for *nanos* mRNA with Me31B-GFP condensates in **(J)** (n = 16). For all plots based on imaging, each data point represents the average colocalization value for all mRNA particles with Me31B-GFP in an image. Significance was assessed using Mann-Whitney statistical tests. Error bars represent standard deviation. **** P < .0001.

To test whether the reduction of *bicoid* mRNA resulted from premature translation or degradation, we examined Bicoid-GFP expression at the anterior of the oocyte where *bicoid* transcripts are most concentrated (Fig. 3E). Ectopic Bicoid-GFP expression was not detected in the *tral^RNAi^* background, indicating that the decrease in *bicoid* mRNA was due to degradation, without translation (Fig. 3F). Similarly, Nanos-GFP was not ectopically expressed during mid-oogenesis (Fig. S3D). Importantly, transcription site activity for neither *bicoid* nor *nanos* mRNA was altered in the absence of Tral (Fig. S3F-H), further supporting a post-transcriptional mechanism of regulation.

Previous work has shown that the physical state of P-bodies is critical for mRNA storage, particularly in oocytes. In late-stage oocytes, 1,6-hexanediol disrupts solid-like P-bodies and releases *bicoid* mRNA, indicating that solid-like condensate properties are essential for transcript retention^11^. However, whether nurse cell P-bodies exhibit similar physical states is unknown. To address this, we treated control nurse cells with low concentrations of 1,6-hexanediol and assessed *bicoid* and *nanos* mRNA colocalization with Me31B-GFP and Cup-YFP. Strikingly, neither mRNA exhibited significant changes in colocalization, suggesting that nurse cell P-bodies are more liquid-like and permissive to dynamic mRNA incorporation (Figs. 3G-I, and S3B, C, E). Treatment with 0.5% SDS, which also disrupts hydrophobic interactions, yielded comparable results (Fig. S3B, C, F).

Given our previous finding that Tral affects intramolecular interactions in P-bodies (Fig. 2) and that Tral is an RBP, we next asked whether the reduced level of *bicoid* mRNA association with P-bodies in *tral^RNAi^*egg chambers resulted from altered condensate properties of P-bodies or from loss of direct Tral:mRNA interactions. To test this, we treated *tral^RNAi^* egg chambers with low-dose 1,6-hexanediol and reassessed mRNA colocalization. Remarkably, *bicoid* mRNA association with Me31B-GFP labeled condensates increased to control levels, while *nanos* mRNA colocalization with P-bodies remained unchanged (Fig. 3G-I). Similar rescue effects were observed with Cup-YFP and following SDS treatment (Fig. S3B, C and E, F). These findings suggest that Tral direct binding is not required for *bicoid* mRNA association with P-bodies, but instead, Tral is necessary to maintain a condensate state conducive to *bicoid* mRNA recruitment/retention.

We next assessed the role of electrostatic interactions in mRNA recruitment/retention by treating control egg chambers with 200mM NaCl. This resulted in decreased colocalization of both *bicoid* (∼49%) and *nanos* (∼26%) mRNAs with Me31B-GFP labeled P-bodies (Fig. 3J-L), a trend recapitulated in Cup-YFP labeled condensates (Fig. S3B, C, G). In contrast, in *tral^RNAi^* egg chambers, there was a ∼30% decrease in *bicoid* mRNA colocalization with Me31B-GFP labeled P-bodies after treatment with NaCl, and there was no effect with Cup-YFP labeled P-bodies (Figs. 3K and S3B). Meanwhile, *nanos* mRNA association with both P-body markers was strongly reduced by NaCl (∼42% with Me31B-GFP, ∼26% with Cup-YFP) (Figs. 3L and S3C).

Together, this data reveals that electrostatic interactions are more broadly required for mRNA retention in P-bodies, while hydrophobic interactions play a more selective role. In the absence of Tral, when Me31B and Cup demix within P-bodies, disruption of hydrophobic interactions paradoxically restores *bicoid* mRNA recruitment, indicating that Tral plays a role in maintaining condensate organization in a way that enables selective mRNA storage. Conversely, *nanos* mRNA remains associated with P-bodies without Tral and after breakdown of hydrophobic interactions. These findings suggest that different maternal mRNAs rely on distinct physical properties of P-bodies for their localization and stability, offering a potential mechanism by which the translational repression of specific transcripts is selectively controlled during oogenesis.

### Tral regulates Me31B and Cup at the transcriptional level

Many P-body proteins have been found to regulate one another, enabling cross-regulatory interactions within the P-body network^16,42^. Previous studies have shown that in a Tral mutant background, Me31B protein and mRNA levels decreased^42^. Consistent with these findings, we observed similar results in our *tral^RNAi^* background where *me31B* mRNA decreased by ∼54% and Me31B protein decreased by ∼63% (Fig. 4A, B). For Cup, we found that mRNA levels increased by ∼35% and Cup protein increased by 383% (Fig. 4A, B). This protein increase was larger than expected based on our imaging data. We suspect this is due to Cup adopting a more solid-like and stable state at higher concentrations in the Tral knockdown background. Notably, the level of *ATP synthase* mRNA, a housekeeping gene, was unaffected, suggesting that these are gene specific changes and not a result of a global effect (Fig. 4A).

**Figure 4:**
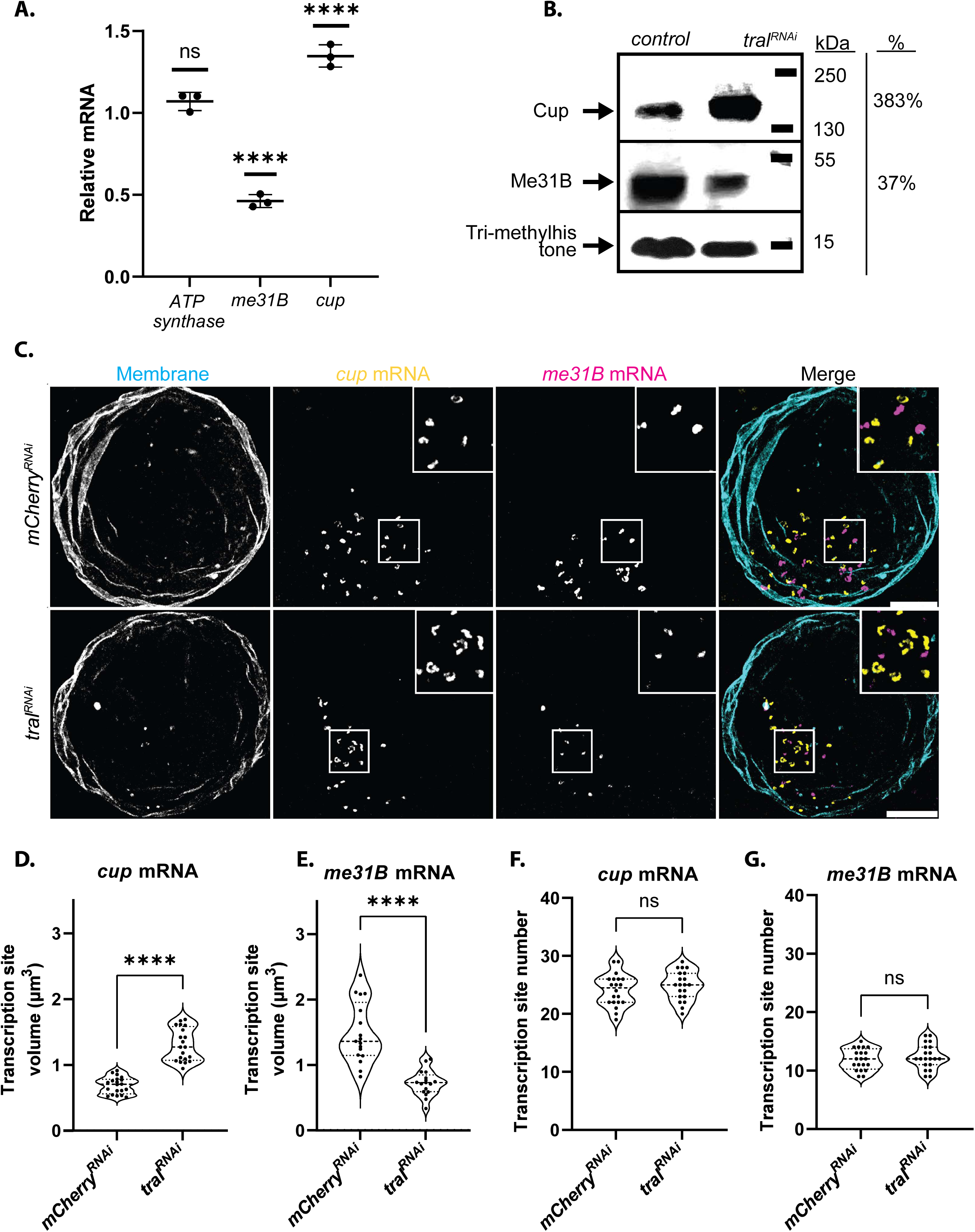
Tral regulates Me31B and Cup at the transcriptional level. **(A)** *ATP synthase, me31B,* and *cup* mRNA levels calculated with RT-qPCR in *tral^RNAi^* egg chambers. Significance calculated with a Welch’s t-test (n = 3). **(B)** Western blot analysis of Cup and Me31B (Tri-methyl-Histone -- loading control). **(C)** Covisualization of smFISH probes labeled *cup* and *me31B* mRNAs with a wheat agglutinin membrane stain labeling the nuclear membrane in *mCherry^RNAi^* and *tral^RNAi^* egg chambers. Images are XY projections of 15 optical Z slices of 0.3µm. Scale bars are 10µm. **(D)** Volume quantification of active *cup* mRNA transcription sites in *mCherry^RNAi^* and *tral^RNAi^* egg chambers (n = 20). **(E)** Volume quantification of active *me31B* mRNA transcription sites in *mCherry^RNAi^* and *tral^RNAi^* egg chambers (n = 17). **(F)** Average *cup* transcription site number per nuclei in *mCherry^RNAi^* and *tral^RNAi^* egg chambers (n = 20). **(G)** Average *me31B* transcription site number per nuclei in *mCherry^RNAi^* and *tral^RNAi^*egg chambers (n = 17). For all plots based on imaging, each data point represents the average value for all transcription sites in an image. Significance was assessed using Mann-Whitney statistical tests. Error bars represent standard deviation. **** P < .0001.

To determine if this differential regulation was occurring at the transcriptional or post-transcriptional level, we next quantified *cup* and *me31B* transcription sites, as the size and the intensity of the sites can indicate transcription rate^43^. Nurse cell nuclei vary in size and DNA content depending on which of the four mitotic divisions they are derived from. In order to remain consistent in our quantitative analysis, we only imaged the nurse cell nuclei that were most proximate to the oocyte nucleus at the dorsal-anterior of the egg chamber^44–46^. *D. melanogaster* nurse cell nuclei are polytene containing many copies of the genome within one nucleus resulting in multiple transcription sites for each gene. Interestingly, in *mCherry^RNAi^*, *cup* exhibited more active transcription sites than *me31B* (∼24 compared to ∼12) despite being located on the same chromosome and having the same copy number (Fig. S4A). *cup* transcription sites were also smaller in size compared to those of *me31B* (Fig. S4B). These findings indicate that the two genes may utilize different strategies for transcriptional regulation; *cup* employs a greater number of transcription sites, which are less active, while *me31B* utilizes fewer sites that are more actively transcribed. We verified that our mRNA nuclear puncta were in fact transcription sites by visualizing their association with DNA (Fig. S4C).

Surprisingly, upon knockdown of Tral, we observed significant changes in the size and intensity of transcription sites. *cup* mRNA nuclear puncta volume increased by ∼89% (from 0.688μm to 2.540μm) and signal intensity increased by ∼47% (Figs. 4C, D and S4D), while *me31B* mRNA puncta displayed the opposite phenotype, becoming ∼51% smaller (from 1.485μm to 0.724μm) and ∼59% less intense (Figs. 4C, E and S4E). Interestingly, despite the observed changes in the volume and intensity of transcription sites, we detected no significant change in the average number of transcription sites compared to control for both genes (Fig. 4F, G). To confirm that these changes in P-body protein transcription sites were specific to *cup* and *me31B,* and not a result of global alterations in transcription, we quantified the transcription sites of maternal mRNAs *bicoid* and *nanos* in nuclei of control and *tral^RNAi^* egg chambers (Fig. S4F). Remarkably, the transcription site volumes of both mRNAs were unaffected by the Tral knockdown (Fig. S4G, H). Together, these findings indicate that Tral specifically plays a role in the transcription of *cup* and *me31B* mRNAs, but not in the availability of their transcription sites.

### Twinstar over-expression partially rescues *me31B* and *cup* transcription levels in the absence of Tral

The observation that *me31B* and *cup* transcription sites are differentially affected by the loss of Tral was particularly striking. Tral knockdown led to the upregulation of one mRNA (*cup*), while downregulating another (*me31B*). To identify potential pathways that could explain this differential transcriptional regulation, we surveyed the literature and found nuclear actin to be a compelling candidate. Actin performs many functions in the nucleus including promoting and repressing transcription depending on gene context^26,47^.

Actin’s structure is tightly regulated by actin-regulatory proteins, with F-actin (fibrous actin) being the predominant functional form in the cytoplasm. F-actin can undergo dynamic remodeling called ‘treadmilling’, where it is polymerized on one end by Chickadee (mammalian homolog: Profilin) and depolymerized on the other end by Twinstar (mammalian homolog: Cofilin)^48,49^. Actin not incorporated into filaments can exist as G-actin, which enters and exits the nucleus through transport processes that also depend on Twinstar and Chickadee, respectively^50,51^.

To assess whether nuclear actin levels were affected by the absence of Tral, we visualized F-actin and G-actin in nurse cells. Overall F-actin organization appeared normal, and no nuclear F-actin fibers were detected even at high laser power that saturated the cytoplasmic signal (Fig. S5A, B). In contrast, quantification of G-actin revealed a decrease in its nuclear enrichment: the nuclear-to-cytoplasmic G-actin intensity ratio was reduced by ∼20% in *tral^RNAi^* egg chambers, indicating lower nuclear G-actin levels in this background (Fig. 5A, B). This reduction was independently confirmed by probing for DNase I, which indirectly labels G-actin, and this gave us similar results (Fig. S5C, D)^52^.

**Figure 5:**
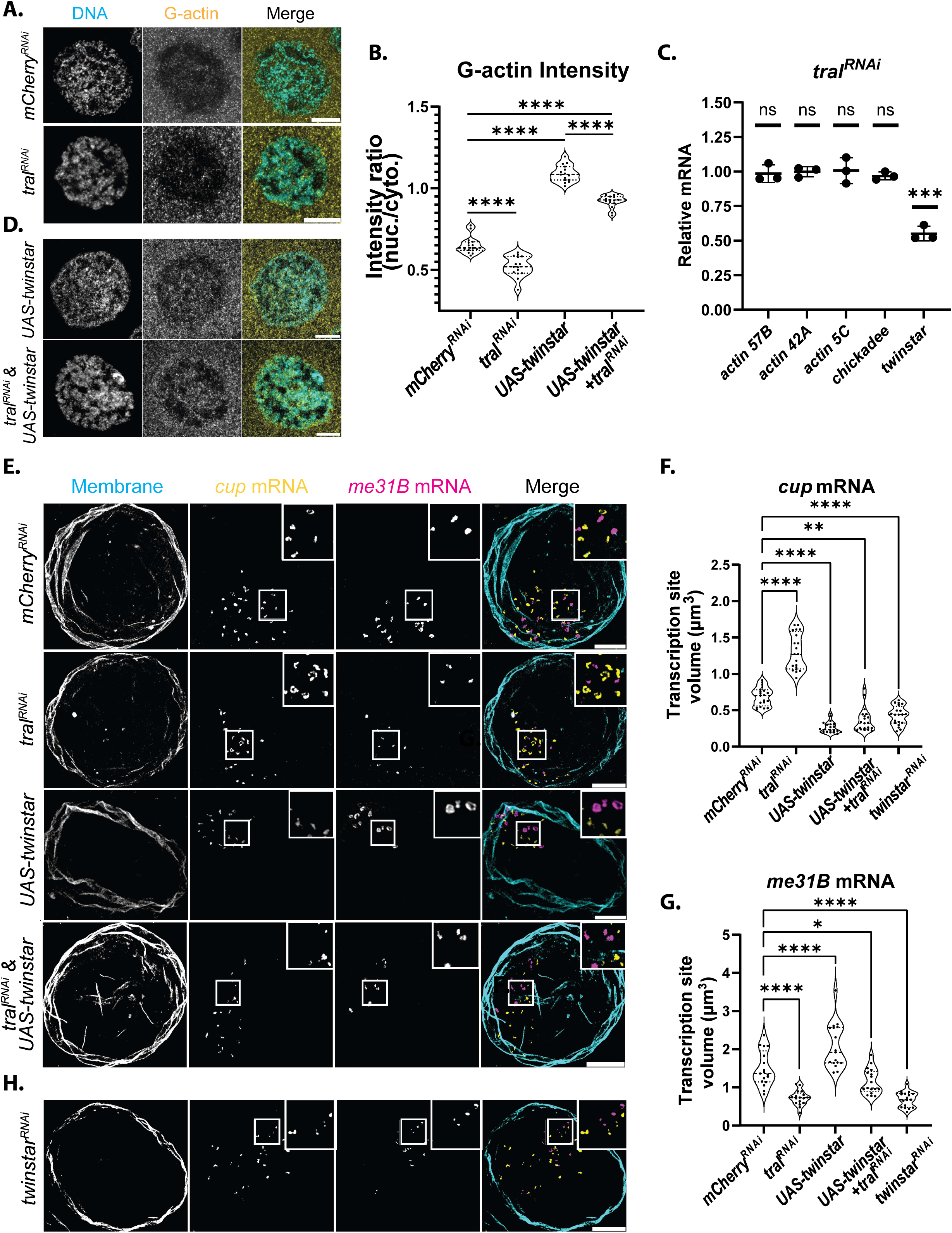
Twinstar over-expression partially rescues *me31B* and *cup* transcription levels in the absence of Tral. **(A)** Visualization of DAPI labeled DNA with immunolabeled G-actin in *mCherry^RNAi^* and *tral^RNAi^* egg chambers. Images are XY projections of 5 optical Z slices of 0.3µm. Scale bars are 10µm. **(B)** G-actin nuclear intensity ratio; calculated by dividing the average G-actin pixel intensity in the nucleus by the average G-actin pixel intensity in the cytoplasm (n = 15). **(C)** *actin 57B, actin 42A, actin 5C, chickadee,* and *twinstar* mRNA levels measured via RT-qPCR in *tral^RNAi^* egg chambers. Significance calculated with Welch’s t-test (n = 3). **(D)** Visualization of DAPI labeled DNA with immunolabeled G-actin in UAS-*Twinstar,* and UAS-*Twinstar* with *tral^RNAi^* egg chambers. Images are XY projections of 5 optical Z slices of 0.3µm. Scale bars are 10µm. **(E)** Covisualization of smFISH probes labeled *cup* and *me31B* mRNAs (wheat agglutinin membrane stain labeling the nuclear membranes) in *mCherry^RNAi^, tral^RNAi^,* UAS-*Twinstar,* and UAS-*Twinstar* with *tral^RNAi^* egg chambers. Images are XY projections of 15 optical Z slices of 0.3µm. Scale bars are 10µm. **(F)** Quantification of active *cup* mRNA transcription sites in *mCherry^RNAi^, tral^RNAi^,* UAS-*Twinstar,* and UAS-*Twinstar* with *tral^RNAi^* egg chambers (n = 20). **(G)** Quantification of active *me31B* mRNA transcription sites in *mCherry^RNAi^, tral^RNAi^,* UAS-*Twinstar,* and UAS-*Twinstar* with *tral^RNAi^*egg chambers (n = 17). **(H)** Covisualization of smFISH labeled *cup* and *me31B* mRNAs (wheat agglutinin membrane stain labeling the nuclear membrane) in *twinstar^RNAi^*egg chambers. Images are XY projections of 15 optical Z slices of 0.3µm. Scale bar is 10µm. For all plots, each data point represents the average value determined per image. Significance was assessed using Mann-Whitney statistical tests. Error bars represent standard deviation. **** P < .0001.

As Tral is an RBP and nuclear actin levels appeared disrupted in the *tral^RNAi^* background, we examined the transcript levels of actin and actin-regulatory proteins in these egg chambers. The actin isoforms predominantly expressed during oogenesis (*actin 57B*, *42A*, and *5C*) were unchanged, and *chickadee* mRNA levels were similarly unaffected. However, *twinstar* mRNA levels were reduced by ∼48% (Fig. 5C). Western blot analysis confirmed that this reduction in mRNA resulted in a ∼41% decrease in Twinstar protein (Fig. S5E). Notably, *twinstar* transcription-site volume remained unchanged in the absence of Tral, indicating that Tral does not regulate *twinstar* transcription and instead acts post-transcriptionally (Fig. S5F, G). Given Twinstar’s established role in G-actin nuclear import, its reduced levels could explain the observed decrease in nuclear G-actin (Fig. 5B)^51^.

To determine whether reduced Twinstar protein levels contributed to the observed transcriptional changes in *cup* and *me31B*, we examined their respective transcription sites in a Twinstar overexpression background. We reasoned that, if Tral affects transcriptional changes via Twinstar, its overexpression should elicit a transcriptional phenotype opposite to that observed in *tral^RNAi^* nuclei. The efficiency of the overexpression was confirmed via RT-qPCR and Western blot analysis (Fig. S5H, I). Notably, when we quantified nuclear actin via an actin antibody and DNase I in the overexpression background, nuclear actin levels increased to more than in control nuclei (Figs. 5B, D, and S5C, D). Strikingly, in this background we observed results opposite to those seen in the Tral knockdown background: *cup* transcription sites were ∼62% smaller while *me31B* transcription sites were ∼40% larger (Fig. 5E-G). These findings support the hypothesis that Tral knockdown reduced Twinstar levels, and that this reduction led to altered transcription site volumes of *cup* and *me31B* mRNA nuclear puncta.

To buttress this, we overexpressed Twinstar in a *tral^RNAi^*background to evaluate if this could rescue the transcription site volumes of *cup* and *me31B* (Fig. 5E). Importantly, the overexpression restored *twinstar* protein and mRNA to near wild-type levels and increased the G-actin levels in the nucleus compared to Tral knockdown alone (Fig. S5C, D, I, and J). Remarkably, we observed that this overexpression almost fully rescued *me31B* mRNA transcription site volumes and partially rescued *cup* mRNA transcription site volumes (Fig. 5E-G). These results indicate that Tral regulates *me31B* and *cup* transcription through its effect on *twinstar* mRNA which leads to a decrease in Twinstar protein. To confirm that the observed changes in transcription site volume correlated with alterations in overall mRNA levels, we assessed global *cup* and *me31B* transcript levels in each background using RT-qPCR. Our results showed that larger transcription sites corresponded to increased overall mRNA levels, while smaller sites resulted in reduced mRNA levels for both *cup* and *me31B* (Fig. S5K, L).

To determine whether reduction in Twinstar, rather than the resulting decreased nuclear actin levels, was responsible for the observed phenotype, we examined *me31B* and *cup* mRNA transcription sites in a Twinstar knockdown background to assess whether they phenocopied the effect of the Tral knockdown on transcription (Fig. 5H). The efficiency of the knockdown was confirmed via RT-qPCR (Fig. S5M). Notably, nuclear G-actin levels decreased lower than in the Tral knockdown background (Fig. S5C, D). We found that, while *me31B* puncta volume decreased in both the *tral^RNAi^* and *twinstar^RNAi^* backgrounds (decreased by ∼51% and ∼54%, respectively), *cup* puncta volume increased in the Tral knockdown, but decreased in the Twinstar knockdown (increased by ∼89% and decreased by ∼38%) (Fig. 5F, G). This result was unexpected; however, previous studies have shown that knockout of the Twinstar homolog, Cofilin, leads to global transcriptional downregulation^53^. To determine whether Twinstar knockdown elicits a similar effect, we assessed transcription of *ATP synthase* and observed an approximate 55% decrease in the volume of its transcription sites (Fig. S5 N, O). These findings indicate that the knockdown of Twinstar may result in a widespread downregulation of transcriptional activity, consistent with previous observations made for Cofilin. Importantly, the opposing behavior of *cup* in Tral versus Twinstar knockdown indicates that its upregulation is not a direct consequence of reduced Twinstar protein. Instead, it may arise from nuclear actin levels being reduced to an intermediate range in Tral-depleted nuclei—a threshold that supports baseline transcriptional activity yet selectively modulates transcription of responsive genes such as *me31B* and *cup*.

### Tral contributes to the organization of *twinstar* mRNA within P-bodies

Given the observed correlation between Tral protein levels and *twinstar* mRNA abundance, we next sought to investigate the mechanism by which Tral regulates *twinstar* mRNA. As Tral is a P-body component and many of its target mRNAs accumulate in P-bodies, we first asked whether *twinstar* mRNA also localizes into P-bodies^54^. To address this, we generated smFISH probes using TFOFinder to visualize *twinstar* mRNA in egg chambers expressing endogenously tagged Me31B-GFP and Cup-YFP (Fig. 6A)^55,56^.

**Figure 6:**
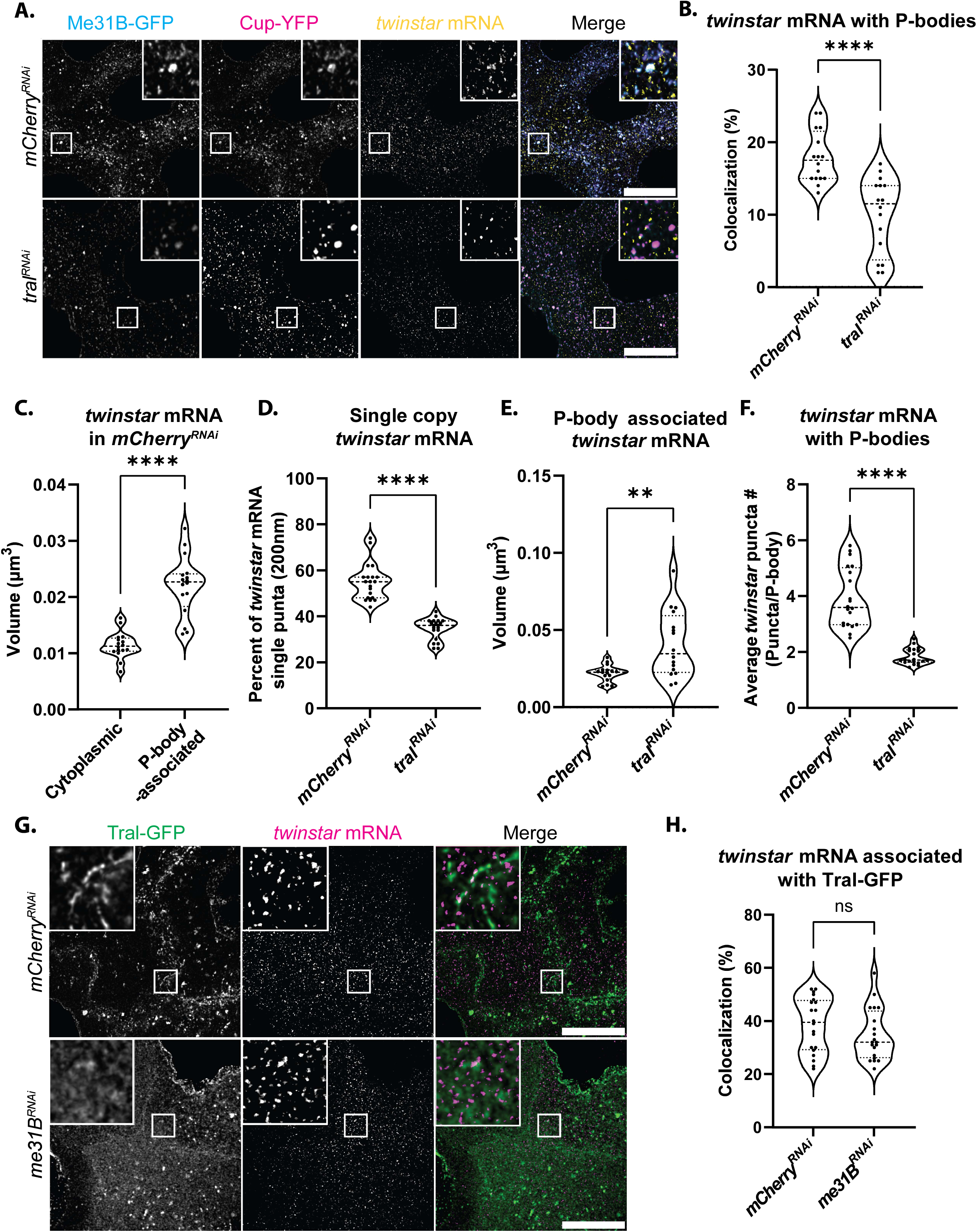
Tral contributes to the organization of *twinstar* mRNA within P-bodies. **(A)** Covisualization of endogenous Me31B-GFP, Cup-YFP, and smFISH probes labeled *twinstar* mRNA in *mCherry^RNAi^* and *tral^RNAi^* egg chambers. Images are XY projections of 5 optical Z slices of 0.3µm. Scale bars are 20µm. **(B)** Colocalization analysis of *twinstar* mRNA with Me31B-GFP:Cup-YFP labeled P-bodies in *mCherry^RNAi^* and *tral^RNAi^* egg chambers (n = 16). **(C)** Volume quantifications comparing cytoplasmic and P-body-associated *twinstar* mRNA puncta in *mCherry^RNAi^* egg chambers (n = 16). **(D)** Percent of *twinstar* mRNA puncta at the diffraction limit (200nm) in *mCherry^RNAi^* and *tral^RNAi^* egg chambers (n = 20). **(E)** Volume quantifications of P-body-associated *twinstar* mRNA puncta in *mCherry^RNAi^* and *tral^RNAi^* egg chambers (n = 16). **(F)** Average *twinstar* mRNA puncta per P-body in *mCherry^RNAi^* and *tral^RNAi^* egg chambers (n = 20).**(G)** Covisualization of endogenous Tral-GFP and smFISH labeled *twinstar* mRNA in *mCherry^RNAi^* and *me31B^RNAi^* egg chambers. Images are XY projections of 5 optical Z slices of 0.3µm. Scale bars are 20µm. **(H)** Colocalization analysis of Tral-GFP with *twinstar* mRNA in *mCherry^RNAi^* and *me31B^RNAi^* egg chambers (n = 20). For all plots, each data point represents the average value determined per image. Significance was assessed using Mann-Whitney statistical tests. Error bars represent standard deviation. **** P < .0001.

We observed that ∼18% of *twinstar* mRNA colocalized with P-bodies, and that these associated puncta were ∼48% larger than those found in the cytoplasm, suggesting that they may contain multiple mRNA molecules (Fig. 4B, C). To examine whether Tral facilitates the association of *twinstar* mRNA with P-bodies, we visualized P-bodies and *twinstar* mRNA in *tral^RNAi^* egg chambers (Fig. 6A). In this background, *twinstar* mRNA colocalization with P-bodies was reduced by ∼44%, indicating that Tral is required either for recruiting *twinstar* mRNA to P-bodies or for maintaining its association (Fig. 6B). Interestingly, we observed an increase in overall *twinstar* mRNA puncta volume in the absence of Tral (Fig. S6A). This appeared to result from a loss of single-copy *twinstar* mRNA, as the number of diffraction-limited puncta (∼200 nm), likely representing individual transcripts, decreased by 38% in *tral^RNAi^* egg chambers (Fig. 6D). Notably, the volume of cytoplasmic *twinstar* mRNA puncta remained unchanged (Fig. S6B), pointing to a specific role for Tral in regulating *twinstar* mRNA within P-bodies.

Further analysis showed that the average size of P-body–associated *twinstar* mRNA puncta increased in the absence of Tral, consistent with the reduction in single-copy species (Fig. 6E). Additionally, the average number of *twinstar* mRNA puncta per P-body decreased by ∼53% (from 3.91 to 1.84), further supporting a role for Tral in organizing *twinstar* mRNA within P-bodies (Fig. 6F).

We next asked whether Tral’s regulation of *twinstar* is dependent on P-body integrity. In *mCherry^RNAi^*egg chambers, ∼39% of *twinstar* mRNA colocalized with Tral-GFP. Interestingly, these puncta were also enriched at the periphery of Tral-GFP marked P-bodies, consistent with previous observations that mRNAs can localize to the edge of P-bodies to facilitate regulated translation^57,58^. Upon Me31B knockdown which disrupts P-body integrity, *twinstar* mRNA:Tral colocalization was not significantly altered (Fig. 6G, H). These findings reveal that Tral promotes the spatial organization of *twinstar* mRNA within P-bodies while also maintaining interactions independent of P-body integrity, underscoring its central role in coordinating mRNA localization and regulation.

### *twinstar* mRNA localization to P-bodies is dependent on association with Tral

As *twinstar* mRNA association with P-bodies was dependent on Tral, we next investigated whether Tral:RNA interactions were required for organizing *twinstar* mRNA within P-bodies. To address this, we chemically disrupted key intramolecular forces in control *mCherry^RNAi^*egg chambers to assess whether chemical treatments could phenocopy the *twinstar* mRNA misorganization phenotype seen in the Tral knockdown egg chambers where *twinstar* mRNA coalesces into fewer, larger puncta withing P-bodies.

We first employed 1,6-hexanediol, as we had previously noted that this rescued *bicoid* mRNA association with P-bodies in the absence of Tral (Fig. 3G, H). Notably, this treatment had no significant effect on the number of *twinstar* mRNA puncta per P-body or on the average volume of P-body–associated *twinstar* mRNA (Fig. 7A-C). Similar results were obtained in Cup-YFP expressing egg chambers (Fig. S7A, B), indicating that hydrophobic interactions are not essential for *twinstar* mRNA organization within P-bodies.

**Figure 7:**
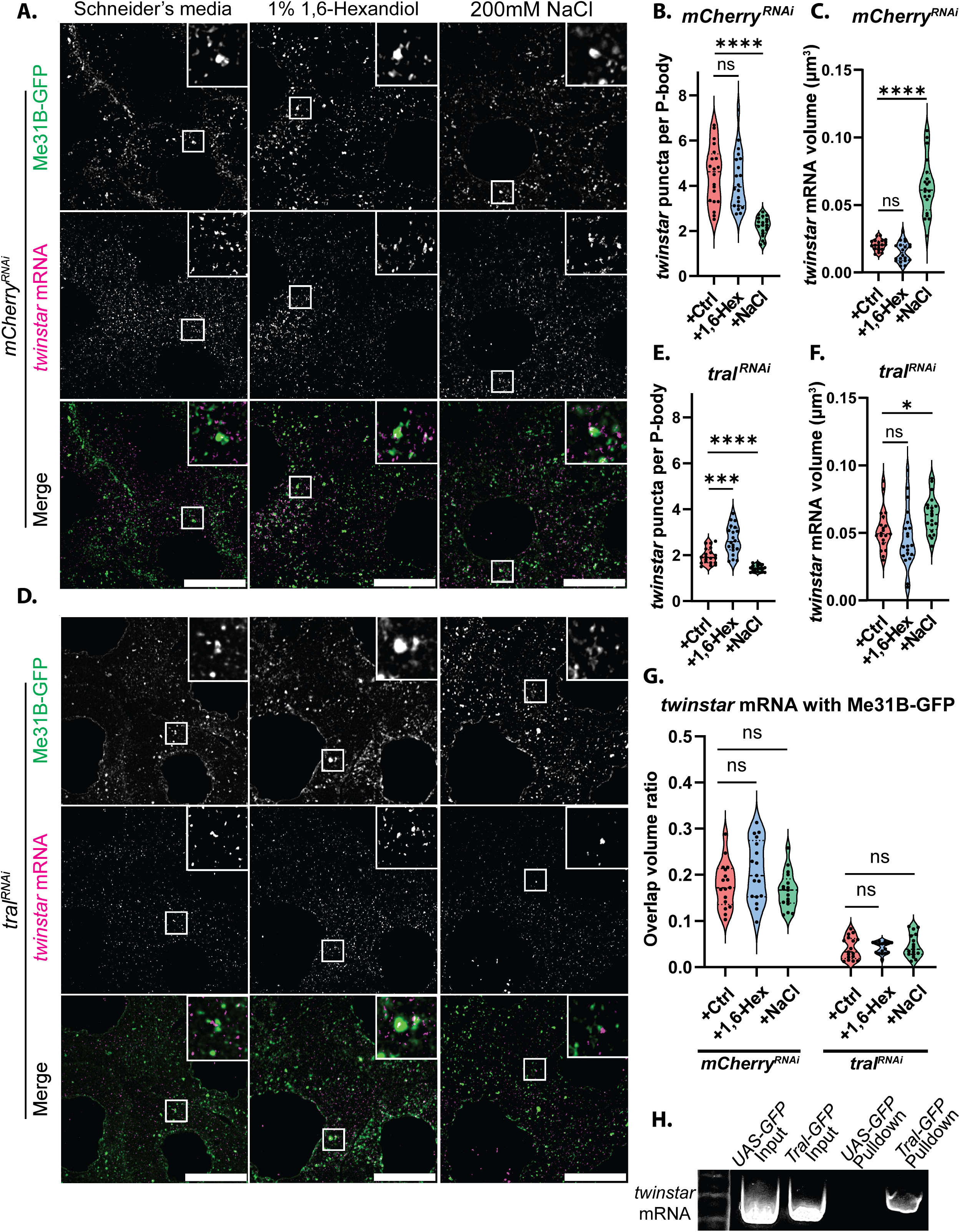
*twinstar* mRNA localization to P-bodies is dependent on association with Tral. **(A)** Me31B-GFP and smFISH labeled *twinstar* mRNA visualized in *mCherry^RNAi^* egg chambers treated with Schneider’s media (+Ctrl), 1% 1,6-hexanediol, or 200mM NaCl. Images are XY projections of 5 optical Z slices of 0.3µm. Scale bars are 20µm. **(B)** Quantification of the average number of *twinstar* mRNA puncta per P-body in each condition in **(A)** (n = 20). **(C)** Calculated volume of the average P-body-associated *twinstar* mRNA puncta in **(A)** (n = 20). **(D)** Me31B-GFP and smFISH labeled *twinstar* mRNA visualized in *tral^RNAi^* egg chambers treated with Schneider’s media (+Ctrl), 1% 1,6-hexanediol, or 200mM NaCl. Images are XY projections of 5 optical Z slices of 0.3µm. Scale bars are 20µm. **(E)** Quantification of the average number of *twinstar* mRNA puncta per P-body in each condition in **(D)** (n = 20). **(F)** Calculated volume of the average P-body-associated *twinstar* mRNA puncta in **(D)** (n = 20). **(G)** Colocalization analysis of *twinstar* mRNA puncta with Me31B-GFP labeled P-bodies across conditions in *mCherry^RNAi^* and *tral^RNAi^* egg chambers. **(H)** UAS-GFP and Tral-GFP RNA pulldowns of *twinstar* mRNA. For all plots, each data point represents the average value detected in an image. Significance was assessed using Mann-Whitney statistical tests. Error bars represent standard deviation. **** P < .0001.

In contrast, treatment with 200mM NaCl for 30 mins, which disrupts electrostatic interactions, produced a marked ∼51% reduction in the number of *twinstar* mRNA puncta per P-body (from 4.48 to 2.20) in Me31B-GFP labeled condensates (Fig. 7B). Intriguingly, while puncta number decreased, the average volume of the remaining *twinstar* mRNA puncta increased by ∼212%, suggesting that electrostatic disruption promotes aggregation of individual transcripts into fewer, larger assemblies (Fig. 7C). A similar trend was observed in Cup-YFP labeled P-bodies, with a ∼63% reduction in puncta number per P-body (from 4.56 to 1.66) and a ∼213% increase in volume (Fig. S7A-C). These results phenocopied the Tral knockdown condition, implicating electrostatic interactions, potentially mediated by Tral, as key to maintaining *twinstar* mRNA organization within P-bodies.

To further explore the role of Tral in this process, we repeated these treatments in the *tral^RNAi^* background. Surprisingly, 1,6-hexanediol treatment increased the number of *twinstar* mRNA puncta per P-body by ∼36% (1.95 to 2.66), without significantly affecting puncta volume (Fig. 7D-F). A comparable ∼21% increase in puncta number was observed in Cup-YFP expressing egg chambers, accompanied by a slight reduction in volume (Fig. S7D-F). These findings suggest that weakening hydrophobic interactions may partially compensate for Tral loss, possibly by promoting dynamic reorganization of RNA-protein complexes.

Conversely, NaCl treatment in the *tral^RNAi^* background caused a ∼28% decrease in puncta number per P-body (from 1.95 to 1.41) and a ∼22% increase in puncta volume in Me31B-GFP marked P-bodies (Fig. 7D-F). Cup-YFP labeled P-bodies showed a similar reduction in puncta number (∼18%, from 1.56 to 1.28) without a significant change in volume (Fig. S7D-F). These results reinforce the idea that electrostatic interactions play a dominant role in organizing *twinstar* mRNA within P-bodies and that Tral contributes to maintaining these interactions.

Given that both Tral knockdown and electrostatic disruption via NaCl reduce *twinstar* mRNA puncta per P-body, we next asked whether the observed loss of organization could also explain the reduced colocalization of *twinstar* mRNA with P-bodies in *tral^RNAi^* egg chambers (Fig. 7B). To test this, we quantified the overlap volume ratio between *twinstar* mRNA and Me31B-GFP or Cup-YFP under each treatment condition. This volumetric approach accounts for differences in puncta size and provides a more accurate representation of true spatial association. Interestingly, we observed no significant differences in overlap volume ratio across conditions (Figs. 7G and S7G). This data suggests that while electrostatic interactions are critical for the internal organization of *twinstar* mRNA within P-bodies, they may not be necessary for its initial association, as overall levels of colocalization remained unchanged.

This observation raised the possibility that Tral may mediate the recruitment of *twinstar* mRNA to P-bodies not by affecting condensate phase properties, but instead by protein:mRNA interactions. Indeed, our earlier data showed a ∼44% reduction in *twinstar* mRNA colocalization with P-bodies in the *tral^RNAi^*background (Fig. 6B). To investigate a possible interaction, we performed a Tral-GFP pulldown assay. Interestingly, we detected enrichment of *twinstar* mRNA in the Tral-GFP pulldown, but not in GFP-only controls, indicating a specific association between Tral and *twinstar* mRNA (Fig. 7H). Together, these findings support a model in which Tral recruits *twinstar* mRNA to P-bodies via protein:mRNA interactions either directly or in a complex and maintains its spatial organization within P-bodies through stabilization of electrostatic interactions.

## DISCUSSION

P-bodies have emerged as critical regulators of mRNA storage. While the role of individual P-body proteins in modulating transcript recruitment to these condensates has been well-documented, the concept that the emergent properties of P-bodies dictate mRNA association is still poorly understood^15,23,28^. In this study, we demonstrate that Tral plays a dual but integrated role in P-body regulation: first, by enforcing a shared condensate state among core P-body proteins, and second, by coupling this condensate organization to transcriptional homeostasis through regulation of *twinstar* mRNA.

This work characterizes a novel mechanism of P-body self-regulation. While previous work has shown that depletion of Tral protein leads to a downregulation of Me31B, we further elucidated the underlying mechanism behind this regulation^42^. We found that Tral is essential for maintaining the stability of *twinstar* mRNA and is responsible for promoting its association with P-bodies. In *tral^RNAi^* egg chambers, *twinstar* mRNA is degraded, and there is a correlative decrease in nuclear actin levels. Whether this reduction in *twinstar* mRNA is due to a shift in P-body function from storage to degradation, or to increased cytoplasmic partitioning of the *twinstar* mRNA which spurs cytoplasmic degradation, is still unknown and warrants further investigation. This reduction coincides with reciprocal changes in *cup* and *me31B* transcription, consistent with prior evidence that nuclear actin can exert gene-specific effects depending on concentration and chromatin context.

We propose that this mechanism may function as a form of P-body autoregulation and, in some instances, serve as a compensatory mechanism for ensuring P-body maintenance. Notably, egg chambers where Me31B is knocked down exhibit lethality at ∼stage 8 of oogenesis. However, when Tral is knocked down, Me31B levels similarly decrease, yet development progresses through oogenesis. This suggests that the upregulation of Cup in the Tral knockdown may compensate to preserve P-body integrity and ensure continued oogenesis. Such genetic compensation is consistent with studies in vertebrates, where *β-actin* knockout induced genomic reprogramming to sustain cell migration^47^.

We previously found that knockdown of Me31B, which disrupts P-body integrity, led to differential regulation of mRNAs: *cyclin A* remains translationally repressed, while *cyclin B* is ectopically released for translation^59^. To further explore the mechanisms underlying transcript-specific regulation within P-bodies, we examined the localization of two maternal mRNAs, *bicoid* and *nanos*. Interestingly, these transcripts were differentially affected by changes in P-body phase state, suggesting that internal biophysical properties of condensates influence transcript specific partitioning. In *tral^RNAi^* egg chambers, *bicoid* mRNA was less associated with P-bodies, but this could be rescued by disrupting hydrophobic interactions, indicating that Tral does not recruit *bicoid* to P-bodies by binding *bicoid* mRNA directly but instead maintains a condensate environment conducive to its recruitment. In contrast, *nanos* mRNA localization was unaffected by Tral depletion, and it associated with P-bodies independent of hydrophobic interactions, with its association relying instead on electrostatic interactions. This finding provides a novel mechanism by which P-bodies could differentially regulate mRNAs within the same condensate.

In the oocyte, some mRNAs must be released from P-bodies, while others require long-term sequestration ^28,40,60,61^. Current models struggle to explain how this selective release and maintenance of transcripts is achieved. Here, we propose that the type of weak interactions mediating mRNA recruitment, electrostatic versus hydrophobic, may act as a biophysical filter: transcripts requiring frequent release may be tethered by weaker, reversible electrostatic interactions, whereas those intended for stable storage are retained through hydrophobic interactions while still others are maintained solely by protein binding. This differential interaction model lends support to the core-shell organization of RNA granules, in which condensate subdomains may differentially regulate transcript behavior^62,63^. Our findings suggest a new conceptual framework in which phase-specific interactions, rather than uniform sequestration, underpin the selective recruitment and release of mRNAs within condensates. This expands the paradigm of P-bodies from passive mRNA storage sites to dynamic regulators of transcript life cycles through phase-tuned interaction specificity.

Here, we also present the first evidence that *twinstar* mRNA is recruited to P-bodies. This finding aligns with previous studies suggesting that P-bodies store mRNAs encoding proteins required for rapid translational activation to facilitate dynamic cellular responses^54^. *twinstar,* a key regulator of actin dynamics, fits this profile, and has been described as a ‘functional node’ in biology because of its ability to receive and transmit cellular information^64^. Twinstar’s homolog, Cofilin, has many roles outside of actin depolymerization and has been shown to function in apoptosis initiation, ER stress response, enzyme activation, as well as acting as a redox sensor^65–67^. Many mRNAs that localize to P-bodies are also known to undergo stable, long-range transport within cells. Notably, *cofilin* mRNA must be transported over long distances within neurons, where it is locally translated to facilitate actin branching in axons^68–70^, and it similarly has been shown to localize to the leading edge of migrating cells to facilitate movement^71^. As such, *twinstar* fits both categories of transcripts typically enriched in P-bodies: those that are involved in maintaining cellular homeostasis and those that require long-distance localization for their function.

Although Tral and Twinstar have not been directly linked in the literature, several lines of evidence suggest they may be functionally connected. Tral has been shown to associate with and be post-transcriptionally regulated by FMRP (Fragile X Mental Retardation Protein), the protein underlying Fragile X Syndrome, and more recently, Cofilin has also been associated with the progression of this disease^72,73^. Furthermore, Cofilin has been identified as a key regulator in the ER stress response, underscoring its role as a critical regulon gene^74,75^. This connection is particularly intriguing given our previous work linking P-body formation with ER exit sites^34^. Taken together, these findings raise the possibility of a functional relationship between Tral and Twinstar, providing a strong rationale for why *twinstar* mRNA is maintained in P-bodies via Tral.

Consistent with this, we found that *twinstar* mRNA localization to P-bodies was significantly reduced in the absence of Tral, and the *twinstar* mRNA that remained, exhibited altered organization, forming larger and fewer puncta. Strikingly, this phenotype could be phenocopied by NaCl treatment, suggesting that Tral may help stabilize electrostatic interactions necessary for proper *twinstar* mRNA partitioning within condensates. These findings support a model in which Tral governs not only the presence of *twinstar* mRNA in P-bodies, but also its spatial organization within them.

We propose a model in which Tral functions as a molecular integrator that links P-body condensate organization to transcriptional homeostasis (Fig. 8). By coordinating hydrophobic and electrostatic interactions between Me31B and Cup, Tral promotes a shared condensate state that supports selective mRNA retention. Within this environment, Tral recruits and organizes *twinstar* mRNA through protein:RNA interactions and stabilization of electrostatic contacts, promoting its storage and stability. Loss of Tral disrupts this organization, leading to reduced *twinstar* mRNA levels, decreased nuclear G-actin, and gene-specific transcriptional changes in core P-body components. These transcriptional shifts, in turn, reshape P-body composition, establishing a potential feedback loop in which condensate material properties, mRNA clients, and gene expression are co-regulated. This framework supports a view of P-bodies as dynamic, phase-tuned regulatory environments rather than passive mRNA storage sites.

**Figure 8:**
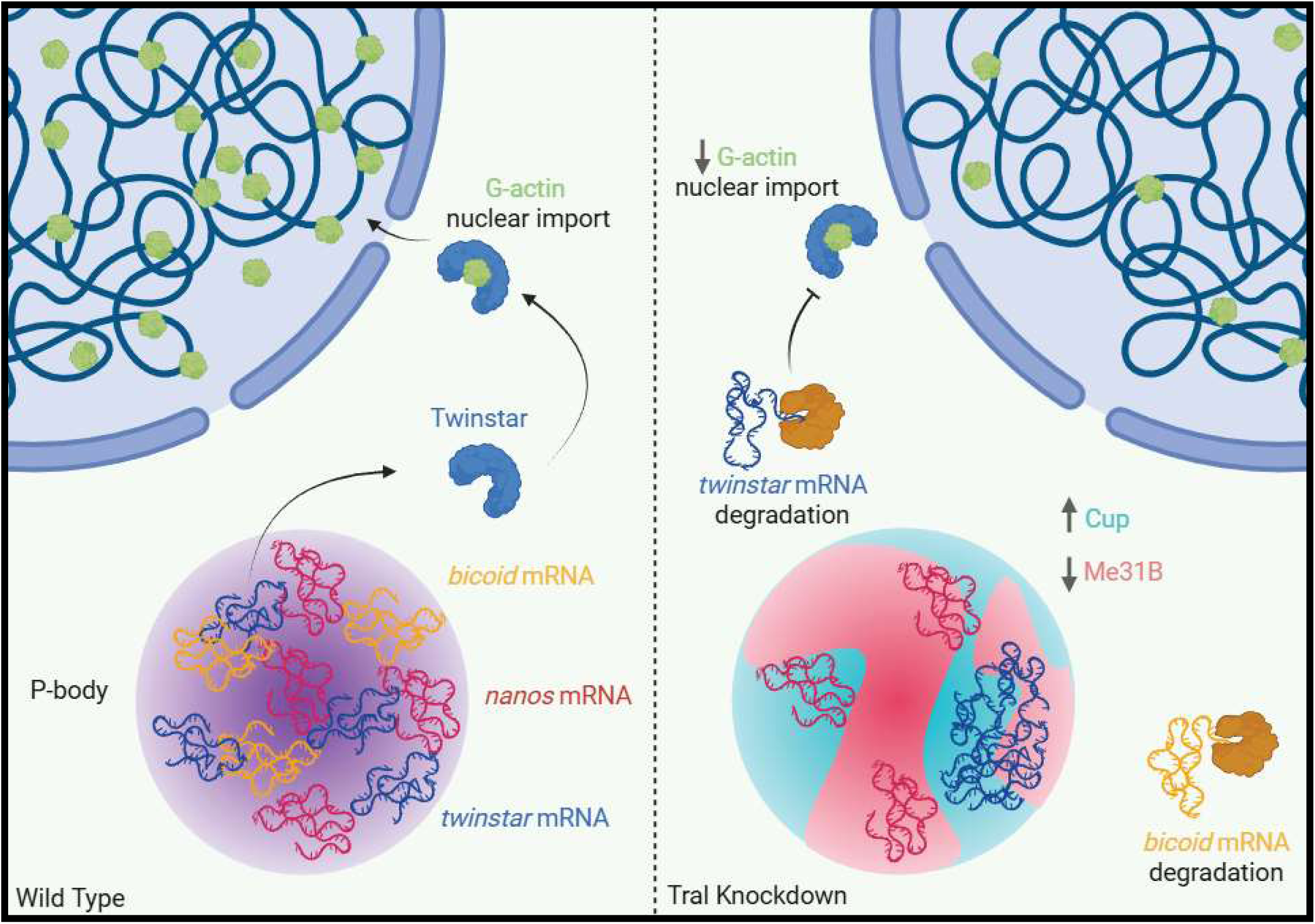
Feedback mechanism coordinated by Tral between P-bodies and *twinstar* mRNA/protein.

## METHODS AND PROTOCOLS

### Fly husbandry

*Drosophila melanogaster* stocks were maintained on standard cornmeal agar food at 25°C. Female flies were put in grape vials and fed yeast paste 2-3 days prior to dissection. Fly stocks obtained from Bloomington *Drosophila* Stock Center: UAS-*mCherry^RNAi^* (BL #35785), UAS-*tral^RNAi^, Me31B-GFP* (BL #51530), *tub-a*(V37)-Gal4 (BL #7063), UAS-*twinstar^RNAi^* (BL #65055), UAS-*Twinstar* (BL #9235), and UAS-GFP (BL #35786). Kyoto *Drosophila* Stock Center: *Cup-YFP* (DGRC 115-161) and Tral-GFP (DGRC 110-584). Bicoid-GFP^76^. *Tral-RFP* was a kind gift from Dr. D. StJohnston (Gurdon Institute at the University of Cambridge), and Nanos-GFP was a kind gift from Dr. E. R. Gavis (Princeton University). UAS-*mCherry^RNAi^* (BL #35785) was used as a control in all RNAi experiments to account for effects of activated RNAi machinery. All RNAi lines were driven by *tub-a*(V37)-Gal4 (BL #7063).

### Immunofluorescence staining of *D. melanogaster* egg chambers

Ovaries were dissected and fixed in 2% PFA in PBS for 10 minutes at room temperature. Fixed egg chambers were washed three times for 10 minutes each in PBST (PBS containing 0.3% Triton X-100), then permeabilized and blocked for 2 hours in PBS containing 1% Triton X-100 and 1% BSA. Samples were incubated with primary antibodies overnight at room temperature with gentle rocking, followed by three 10-minute washes in PBST. Secondary antibody incubation was carried out using fluorescently labeled antibodies (1:1000; DyLight 650; ThermoFisher) for 2 hours at room temperature, followed by three additional 10-minute washes in PBST. Samples were mounted in RapiClear (SUNjin Lab) mixed with Aqua-Poly/Mount (Polysciences) at a 75:25 ratio for imaging. The following antibodies and stains were used: mouse anti-Cup (1:1000), a generous gift from Dr. A. Nakamura (Institute of Molecular Embryology and Genetics, Kumamoto University), mouse anti-actin (clone JLA20, 1:200), deposited to the Developmental Studies Hybridoma Bank (DSHB) by J.J.-C. Lin, and Phalloidin–Alexa Fluor 647 (1:200; Life Technologies) for F-actin staining.

### smFISH labeling

Single-molecule fluorescence *in situ* hybridization (smFISH) was performed following the protocol described by Bayer et al. (2015)^77^, with minor modifications. Ovaries were dissected and fixed in 4% PFA in PBS for 10 minutes at room temperature. Fixed egg chambers were washed three times for 10 minutes each in 2x SSC, then pre-hybridized with a 15-minute wash in 2x SSC containing 10% formamide. Samples were then incubated overnight at 37 °C with smFISH probes (1:50) and a nuclear membrane stain, wheat germ agglutinin conjugated to CF405S (1:50; Biotium). Following hybridization, egg chambers were washed three times for 10 minutes each in pre-warmed 2x SSC with 10% formamide at 37 °C and mounted in ProLong Diamond Antifade Mountant (Life Technologies) for imaging. The following probe sets were used: *nanos* mRNA labeled with 48 Quasar 670 probes, *bicoid* mRNA labeled with 48 Quasar 570 probes, *me31B* mRNA labeled with 30 eGFP recognizing Quasar 670 probes, *cup* mRNA labeled with 48 Cal Fluor Red 590 probes. Probes were made by Biosearch technologies.

*twinstar* mRNA was labeled using 14 Atto 633-conjugated smFISH probes, designed and synthesized following the protocol described by Gaspar et al. (2017)^56^. Probe sequences were computationally determined using TFOFinder^55^. *In situ* hybridization was performed as previously described.

### Chemical treatments

Egg chambers were dissected directly into Schneider’s media alone (+Control) or supplemented with 1% 1,6-Hexanediol, 200mM NaCl, or 0.5% SDS. Following 30 minutes incubation, egg chambers were rinsed with PBS 1X and then fixed and mounted in RapiClear (SUNjin Lab) mixed with Aqua-Poly/Mount (Polysciences) at a 75:25 ratio for imaging as previously described.

### Microscopy

All imaging was performed using a Leica TCS SP8 Laser Scanning Confocal Microscope equipped with a white light laser (470–670 nm), a 405 nm solid-state laser, and a continuous-wave STED 660 nm high-intensity laser. For confocal imaging, a 63x/1.4 NA oil immersion objective was used, and optical Z-sections were acquired at 0.3 µm intervals. For STED imaging, a 100x/1.4 NA oil immersion objective was used with a zoom factor of 5x, and optical Z-sections were acquired at 0.22 µm intervals. STED samples were prepared from 25 µm ovary cryosections. All images were acquired using an automated XYZ piezoelectric stage and saved as 16-bit files. Image acquisition was carried out using Leica LAS X software.

### Tissue preparation for sectioning

Ovaries were dissected directly into 4% PFA in PBS and fixed for 10 minutes at room temperature. Samples were then washed three times for 10 minutes each in PBST (PBS with 0.1% Triton X-100), followed by a 5-minute wash in 0.1 M glycine, pH 3.0. After fixation and washing, ovaries were incubated overnight at 4 °C in 30% sucrose. The following day, samples were embedded in O.C.T. Compound (Tissue-Tek) and flash frozen prior to sectioning. Cryosections of 25µm thickness were prepared using a cryotome and stored at −80 °C until use.

### Imaging analysis

Identical acquisition settings were used for all control and experimental samples to ensure comparability. For each condition, images were obtained from similar stage egg chambers from three independent experiments, each prepared from a separate fly cross. All raw images were deconvolved prior to analysis using Leica’s Lightning deconvolution module. Image processing was carried out using consistent batch parameters across all samples. Quantitative analyses were performed using Imaris Microscopy Image Analysis software (Oxford Instruments). P-bodies, proteins, and mRNAs were detected using the Imaris “Surface” module, which enables object-based segmentation and object-to-object spatial analysis. Colocalization was determined using the “Surface–Surface” analysis function, where objects with a shortest edge-to-edge distance of less than 0 µm were classified as colocalized. All statistical analyses were conducted using Mann–Whitney statistical tests in GraphPad Prism 8 (GraphPad Software). Figures were assembled and image adjustments applied uniformly using Fiji/ImageJ (NIH) ^78^.

### Western blot analysis

For each genotype, ten ovaries were dissected directly into 95 µL of 2x Laemmli Sample Buffer (Bio-Rad) supplemented with 5 µL β-mercaptoethanol (BME) and immediately subjected to mechanical lysis. Samples were heated at 95 °C for 10 minutes and then centrifuged at 10,000xg for 10 minutes at 4 °C. Clarified lysates were loaded onto 10% SDS-PAGE acrylamide gels for electrophoretic separation. Primary antibodies used included mouse anti-Cup (1:3000) and mouse anti-Me31B (1:2000), both generously provided by Dr. A. Nakamura (Institute of Molecular Embryology and Genetics, Kumamoto University); rabbit anti-Tri-methyl-Histone H3 (C42D8) (1:150,000; Cell Signaling Technology); rabbit anti-Bicaudal D (BicD) clone 1B11 (1:100), obtained from the Developmental Studies Hybridoma Bank (DSHB) courtesy of R. Steward; and rabbit anti-Cofilin (10960-1-AP) (1:3000; Proteintech). Bands were detected using TrueBlot ULTRA secondary antibodies (anti-mouse and anti-rabbit IgG HRP, 1:50,000; Rockland) and visualized with SuperSignal West Femto Maximum Sensitivity Substrate (ThermoFisher Scientific).

### RNA isolation and RT-qPCR

Whole ovaries were dissected into ice-cold PBS (4 °C) and immediately subjected to mechanical lysis in TRIzol reagent (ThermoFisher Scientific) for total RNA extraction. RNA was precipitated and washed with ethanol and resuspended in RNase-free water. RNA concentration and purity were assessed prior to downstream applications. Reverse transcription was performed using 2.5µg of total RNA with the Superscript IV First-Strand Synthesis Kit (Life Technologies) according to the manufacturer’s instructions. Primers were designed using the DRSC FlyPrimerBank and synthesized by Integrated DNA Technologies. Quantitative PCR reactions were carried out on a Roche LightCycler 480 system (Roche Molecular Systems, Inc.). Each 10µL reaction contained 1µL of cDNA template, 4µL of 10µM primer mix, and 5µL of SYBR Green I Master Mix (Roche Diagnostics). Reactions were performed in triplicates. Relative expression levels were calculated and normalized using Rp-49, and statistical significance was evaluated using a two-tailed unpaired Welch’s t-test.

### RNA pull-down

Ovaries from flies expressing Tral-GFP or GFP alone (20 ovaries per condition) were dissected and immediately fixed in 4% PFA in PBS for 10 minutes at room temperature. Samples were subsequently washed three times for 10 minutes each in PBST (PBS with 0.3% Triton X-100) and transferred to 200µL of RIPA buffer (50 mM Tris-Cl, pH 7.5; 150 mM NaCl; 1% NP-40; 1 mM EDTA) supplemented with EDTA-free Roche cOmplete protease inhibitor cocktail (Sigma-Aldrich) and RiboLock RNase inhibitor (ThermoFisher Scientific). Ovaries were mechanically lysed, and lysates were clarified by centrifugation at 12,000xg for 10 minutes at 4 °C. Cleared lysates were incubated with 10µL of pre-equilibrated GFP-Trap Dynabeads (Chromotek) for 1 hour at room temperature with gentle inversion. Beads were then washed five times for 10 minutes each with supplemented RIPA buffer. mRNA was eluted from the beads using TRIzol reagent (ThermoFisher Scientific), and RNA isolation was performed as previously described. Reverse transcription was conducted to generate cDNA, followed by PCR amplification using *twinstar*-specific primers (Integrated DNA Technologies). PCR reactions were assembled with 6.5µL cDNA, 6µL of 10µM primers, and 12.5µL of PCR Master Mix (Promega), and amplification was carried out on a Veriti 96-well thermocycler (Applied Biosystems). Amplified DNA products were resolved on a 2% agarose gel stained with SYBR Green I Nucleic Acid Stain (Lonza) and visualized under UV illumination.

## Graphics

BioRender was used to prepare Figures 1A, 7E, and 8.

## ACKNOWLEDGEMENTS

We thank Dr. A. Nakamura, (Riken Center for Developmental Biology) for the kind gifts of antibodies. We extend thanks to Dr. D. St Johnston (University of Cambridge) and Dr. E. R Gavis (Princeton University) for the kind gifts of *D. melanogaster* lines. We thank the BDSC Indiana and DGRC Kyoto for providing *D. melanogaster* lines, as well as the TRiP at Harvard Medical School (NIH/NIGMS RO1-GM084947) for the transgenic RNAi stocks. We thank the Bioimaging Facility at Hunter College for access to the Leica TCS SP8 and Imaris -- Image Analysis Software. We thank Dr. P. Feinstein (Hunter College) for allowing us to use the Roche Light-cycler instrument. We also thank Dr. I. E. Catrina (Yeshiva University) for her kind help in designing *twinstar* mRNA smFISH probes.

## DATA AVAILABILITY

This study includes no data deposited in external repositories.

