## Supplemental Figures for "Trailer Hitch tunes condensate phase behavior and mRNA partitioning to regulate P-body homeostasis during *Drosophila melanogaster* oogenesis"

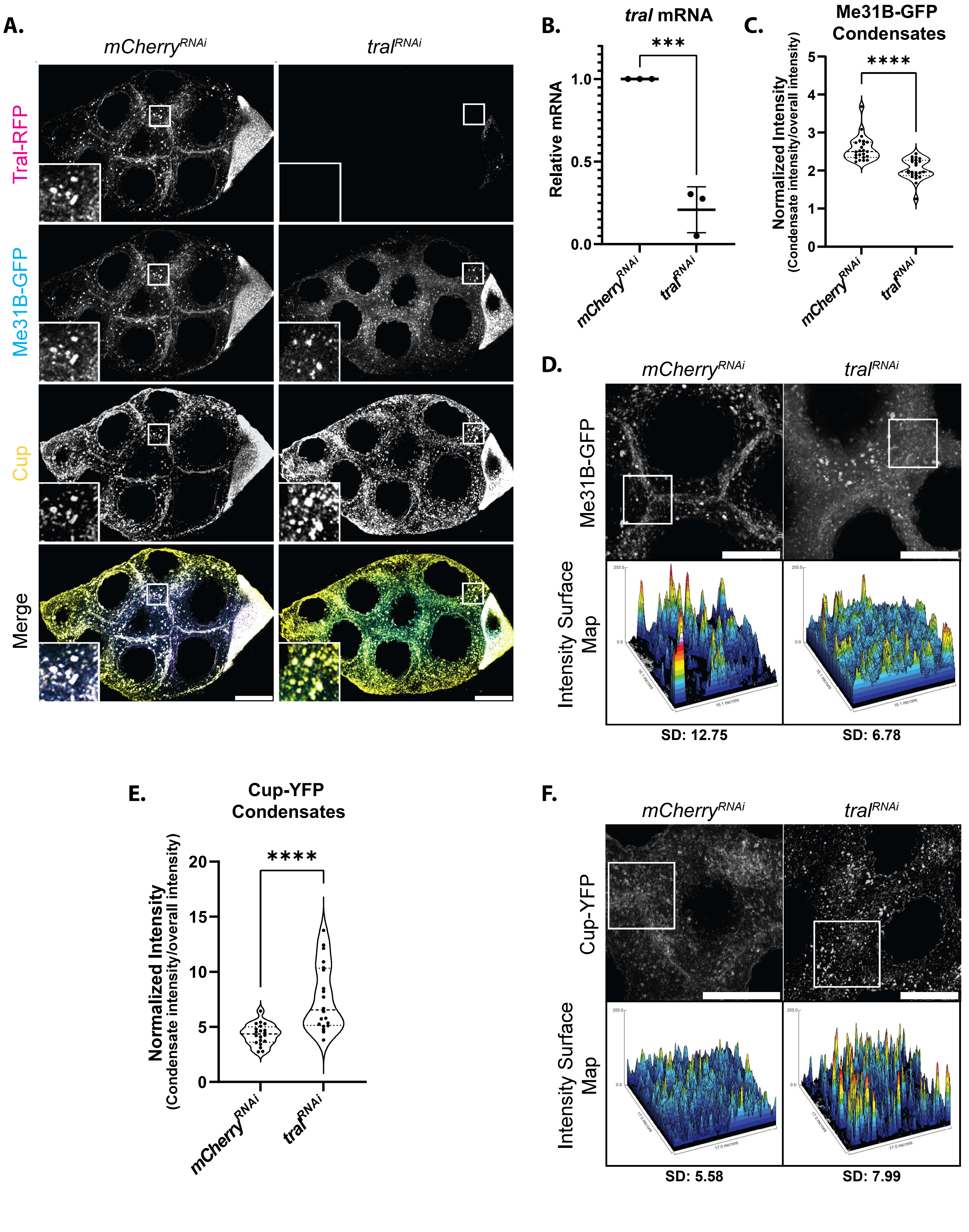


*Fig. S1 (Milano et al, 2026)*

**Figure S1: Tral differentially affects the incorporation of Me31B and Cup into P-bodies.**

**(A)** Covisualization of endogenous Tral-RFP and Me31B-GFP with immunolabeled Cup-YFP in *mCherry*$^{RNAi}$ and *tral*$^{RNAi}$ egg chambers. Images are XY projections of 5 optical Z slices of 0.3µm. Scale bars are 20µm.

**(B)** RT-qPCR analysis of *tral* mRNA in *mCherry*$^{RNAi}$ and *tral*$^{RNAi}$ egg chambers. Significance calculated with Welch's t-test. (n = 3).

**(C)** Normalized fluorescence intensity measurements where puncta fluorescence intensity is divided by the overall image fluorescence intensity for Me31B-GFP labeled condensates (n = 23).

**(D)** 3D intensity maps of Me31B-GFP in selected ROI with calculated standard deviation (SD) of cytoplasmic region fluorescence intensity in *mCherry*$^{RNAi}$ and *tral*$^{RNAi}$ egg chambers**.**

**(E)** Normalized intensity measurements where puncta intensity is divided by the overall image intensity for Cup-YFP labeled condensates (n = 20).

**(F)** 3D intensity maps of Me31B-GFP in selected ROI with calculated standard deviation (SD) of cytoplasmic region fluorescence intensity in *mCherry*$^{RNAi}$ and *tral*$^{RNAi}$ egg chambers.

For all plots based on imaging, each data point represents the average colocalization value for all P-bodies detected in an image. Significance was assessed using Mann-Whitney statistical tests. Error bars represent standard deviation. **** P < .0001.

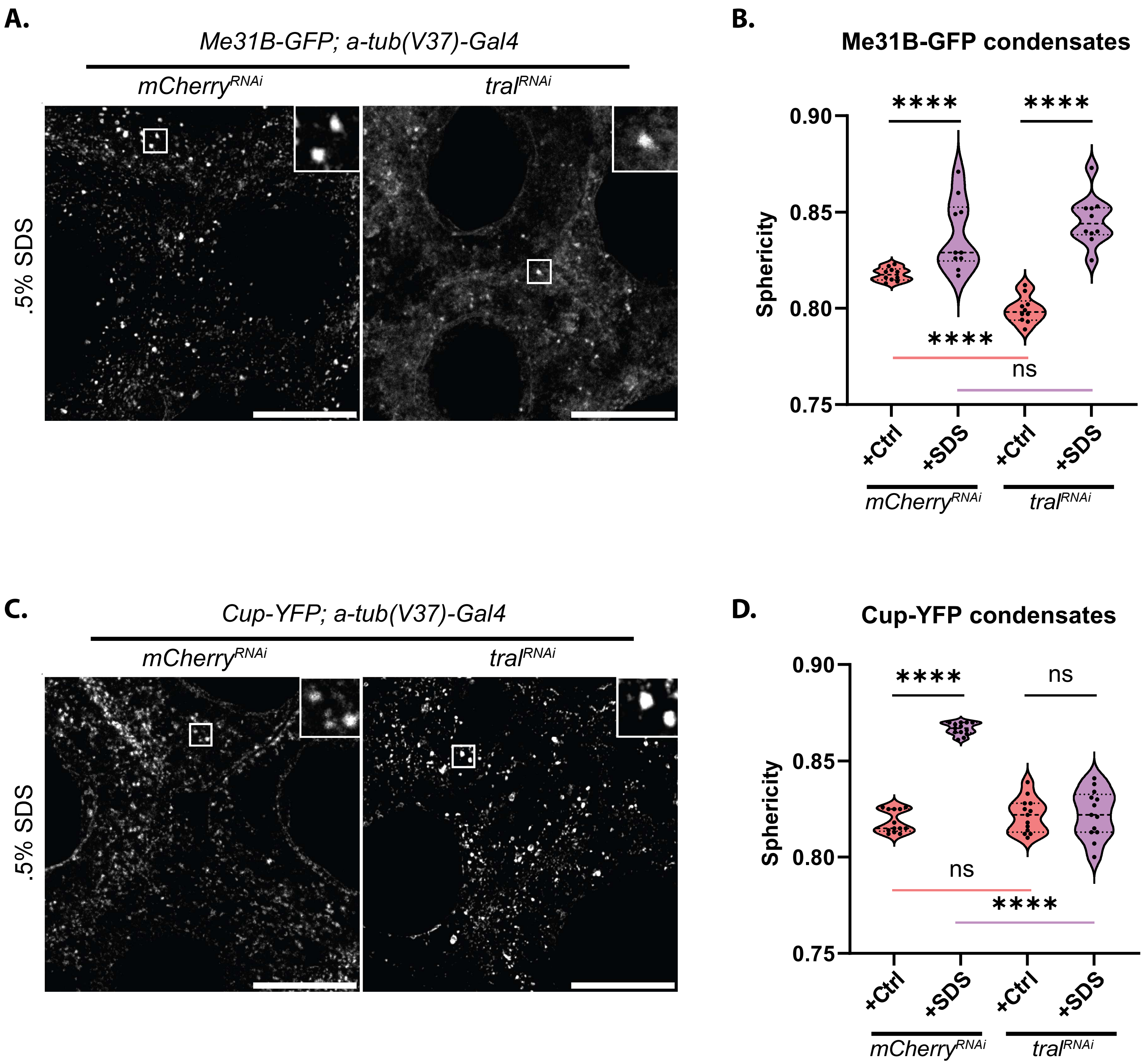


***Fig. S2 (Milano et al, 2026)***

**Figure S2: Tral promotes a shared condensate state between Me31B and Cup.**

**(A)** Me31B-GFP visualized in *mCherry*$^{RNAi}$ and *tral*$^{RNAi}$ egg chambers treated with 0.5% SDS in Schneider's media (+Control). Images are XY projections of 5 optical Z slices of 0.3µm. Scale bars are 20µm.

**(B)** Sphericity quantifications comparing Me31B-GFP labeled condensates in *mCherry*$^{RNAi}$ and *tral*$^{RNAi}$ egg chambers under conditions in **(A)** (n = 10).

**(C)** Cup-YFP visualized in *mCherry*$^{RNAi}$ and *tral*$^{RNAi}$ egg chambers treated with 0.5% SDS in Schneider's media (+Control). Images are XY projections of 5 optical Z slices of 0.3µm. Scale bars are 20µm.

**(D)** Sphericity quantifications comparing Cup-YFP labeled condensates in *mCherry*$^{RNAi}$ and *tral*$^{RNAi}$ egg chambers under conditions in **(C)** (n = 13).

For all plots, each data point represents the average value of all P-bodies detected in an image. Significance was assessed using Mann-Whitney statistical tests. Error bars represent standard deviation. **** $P < .0001$.

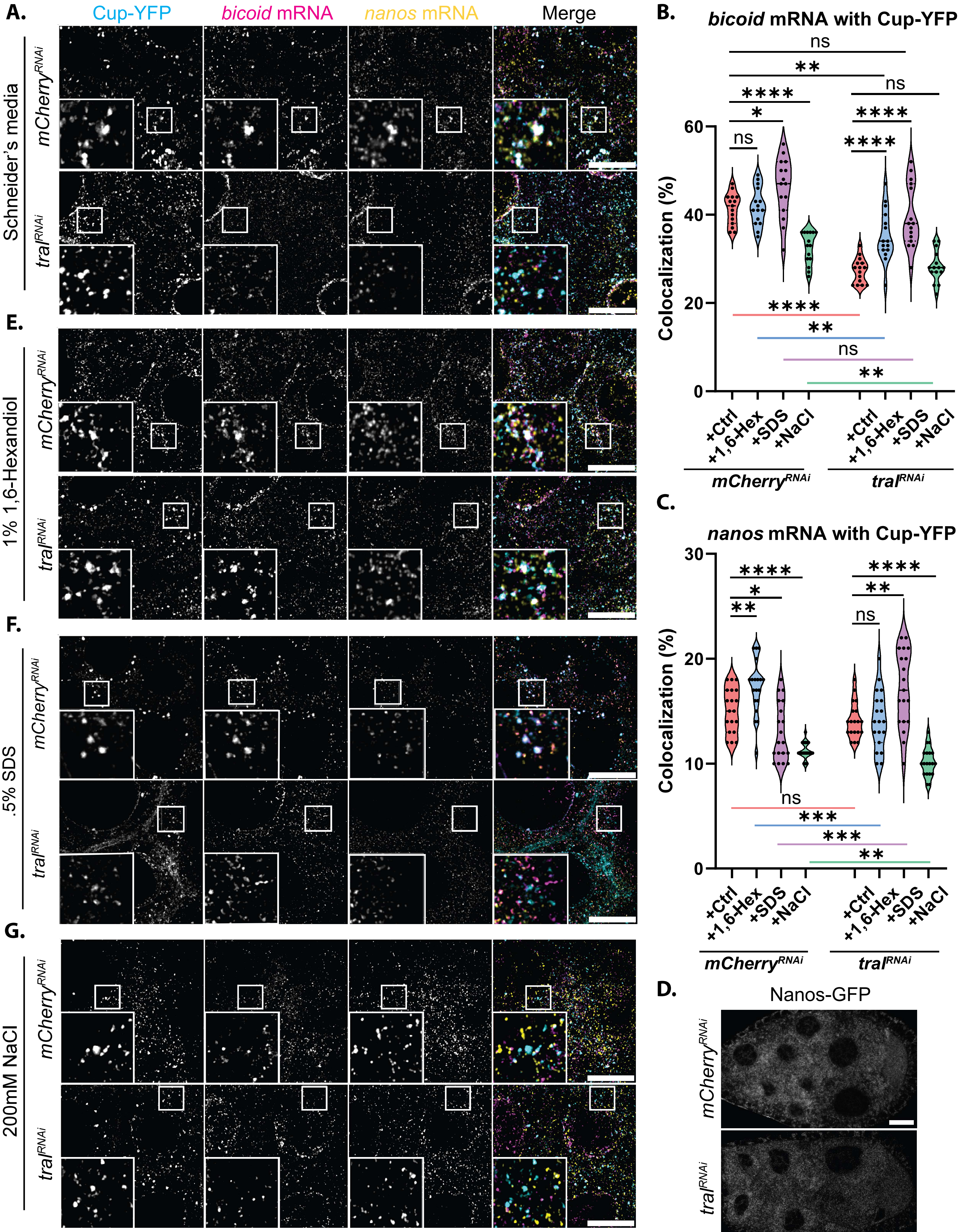


*Fig. S3 (Milano et al, 2026)*

**Figure S3: Tral is necessary for maintaining P-body function in select transcript storage.**

**(A)** Cup-YFP covisualized with smFISH probes labeled *bicoid* and *nanos* mRNAs, in $mCherry^{RNAi}$ and $tral^{RNAi}$ egg chambers incubated in Schneider's media (+Control). Images are XY projections of 5 optical Z slices of 0.3μm. Scale bars are 20μm.

**(B)** Colocalization calculations for *bicoid* mRNA with Me31B-GFP condensates across conditions (**A** and **E-G**) (n = 15).

**(C)** Colocalization calculations for *nanos* mRNA with Me31B-GFP condensates across conditions (**A** and **E-G**) (n = 19).

**(D)** Nanos-GFP visualized in $mCherry^{RNAi}$ and $tral^{RNAi}$ egg chambers. Scale bars are 20μm.

**(E)** Cup-YFP covisualized with smFISH probes labeled *bicoid* and *nanos* mRNAs, in $mCherry^{RNAi}$ and $tral^{RNAi}$ egg chambers incubated in 1% 1,6-hexanediol, **(F)** in 0.5% SDS, and **(G)** in 200mM NaCl. Images are XY projections of 5 optical Z slices of 0.3μm. Scale bars are 20μm.

For all plots, each data point represents the average colocalization value for all mRNA particles with Cup-YFP in an image. Significance was assessed using Mann-Whitney statistical tests. Error bars represent standard deviation. **** P < .0001.

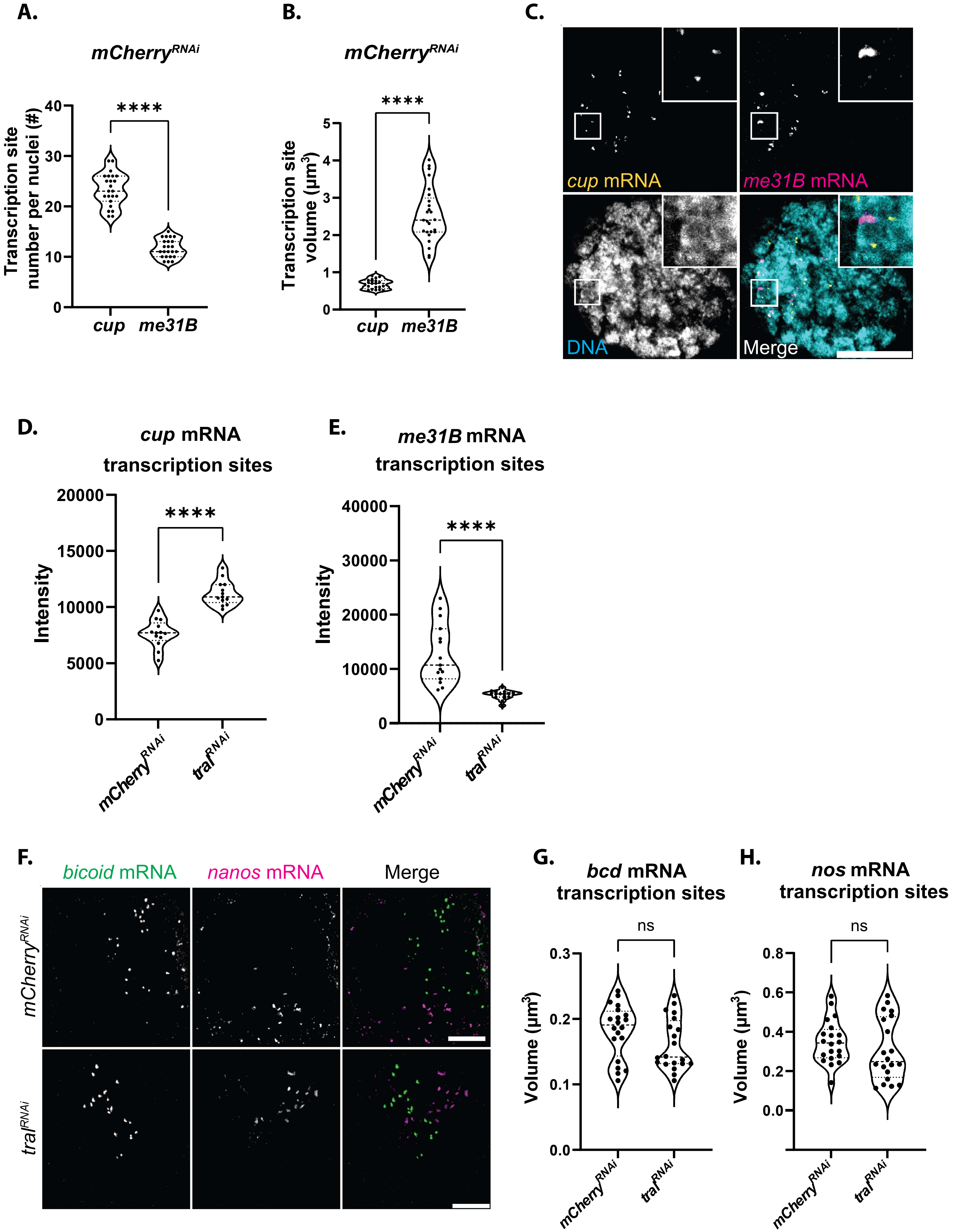


*Fig. S4(Milano et al, 2026)*

**Figure S4: Tral regulates Me31B and Cup at the transcriptional level.**

**(A)** Average transcription site number per nuclei of *cup* and *me31B* in *mCherry*$^{RNAi}$ egg chambers (n = 25).

**(B)** Average transcription site volume of *cup* and *me31B* in *mCherry*$^{RNAi}$ egg chambers (n = 25).

**(C)** Covisualization of DAPI labeled DNA and smFISH probes labeled *cup* and *me31B* transcription sites in *mCherry*$^{RNAi}$ egg chambers. Image is an XY projection of 5 optical Z slices of 0.3µm. Scale bars are 20µm.

**(D)** Average intensity of *cup* mRNA transcription sites in *mCherry*$^{RNAi}$ and *tral*$^{RNAi}$ egg chambers (n = 13).

**(E)** Average intensity of *me31B* mRNA transcription sites in *mCherry*$^{RNAi}$ and *tral*$^{RNAi}$ egg chambers (n = 15).

**(F)** Covisualization of smFISH probes labeled *bicoid* and *nanos* transcription sites in *mCherry*$^{RNAi}$ and *tral*$^{RNAi}$ egg chambers. Images are XY projections of 5 optical Z slices of 0.3µm. Scale bars are 10µm.

**(G)** Average transcription site volume of *bicoid* in *mCherry*$^{RNAi}$ and *tral*$^{RNAi}$ egg chambers (n = 20).

**(H)** Average transcription site volume of *nanos* in *mCherry*$^{RNAi}$ and *tral*$^{RNAi}$ egg chambers (n = 20).

For all plots based on imaging, each data point represents the average value for all transcription sites in an image. Significance was assessed using Mann-Whitney statistical tests. Error bars represent standard deviation. **** P < .0001.

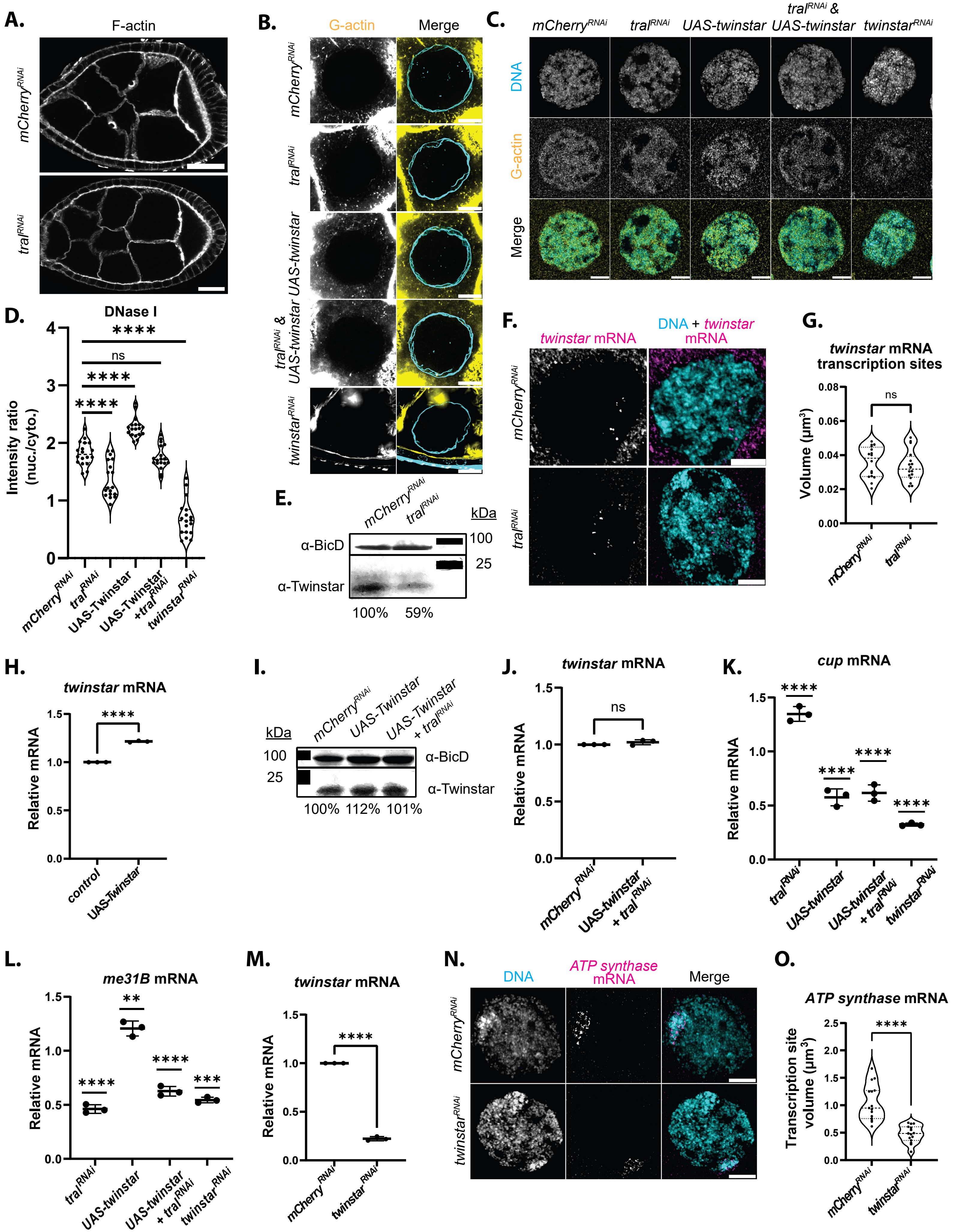


***Fig. S5 (Milano et al, 2026)***

**Figure S5: Twinstar over-expression partially rescues *me31B* and *cup* transcription levels in the absence of Tral.**

**(A)** F-actin visualized by phalloidin staining in control (*mCherry*$^{RNAi}$) and *tral*$^{RNAi}$ egg chambers. Images are XY projections of 5 optical Z slices of 0.3µm. Scale bars are 20µm.

**(B)** Nuclear F-actin visualized by over exposing phalloidin staining in *mCherry*$^{RNAi}$ and *tral*$^{RNAi}$ nurse cell nuclei. Images are XY projections of 5 optical Z slices of 0.3µm. Scale bars are 10µm.

**(C)** Visualization of DAPI labeled DNA with immunolabeled DNase I in *mCherry*$^{RNAi}$, *tral*$^{RNAi}$, UAS-*Twinstar,* UAS-*Twinstar* in *tral*$^{RNAi}$, and *twinstar*$^{RNAi}$ egg chambers. Images are XY projections of 5 optical Z slices of 0.3µm. Scale bars are 10µm.

**(D)** DNase I nuclear intensity ratio; divided the average DNase I pixel fluorescence intensity in the nucleus by the average DNase I pixel fluorescence intensity in the cytoplasm (n = 15).

**(E)** Western blot analysis of Twinstar (BicD -- loading control) in *mCherry*$^{RNAi}$ and *tral*$^{RNAi}$ egg chambers.

**(F)** *twinstar* mRNA visualized with DAPI labeled DNA in *mCherry*$^{RNAi}$ and *tral*$^{RNAi}$ egg chambers.

**(G)** Average *twinstar* mRNA transcription site volume in *mCherry*$^{RNAi}$ and *tral*$^{RNAi}$ egg chambers (n = 16).

**(H)** RT-qPCR analysis of *twinstar* mRNA in control and UAS-*Twinstar* egg chambers. Significance calculated with Welch's t-test. (n = 3).

**(I)** Western blot analysis of Twinstar (BicD -- loading control) in *mCherry*$^{RNAi}$, UAS-*Twinstar*, and UAS-*Twinstar* with *tral*$^{RNAi}$ egg chambers.

**(J)** RT-qPCR analysis of *twinstar* mRNA in *mCherry*$^{RNAi}$ and UAS-*Twinstar* with *tral*$^{RNAi}$ egg chambers. Significance calculated with Welch's t-test (n = 3).

**(K)** RT-qPCR analysis of *cup* mRNA in *tral*$^{RNAi}$, UAS-*Twinstar*, UAS-*Twinstar* with *tral*$^{RNAi}$, and *twinstar*$^{RNAi}$ egg chambers. Significance calculated with Welch's t-test (n = 3).

**(L)** RT-qPCR analysis of *me31B* mRNA in *tral*$^{RNAi}$, UAS-*Twinstar*, UAS-*Twinstar* with *tral*$^{RNAi}$, and *twinstar*$^{RNAi}$ egg chambers. Significance calculated with Welch's t-test (n = 3).

**(M)** RT-qPCR analysis of *twinstar* mRNA in *mCherry*$^{RNAi}$ and *twinstar*$^{RNAi}$ egg chambers. Significance calculated with Welch's t-test (n = 3).

**(N)** *ATP synthase* mRNA visualized with DAPI labeled DNA in *mCherry*$^{RNAi}$ and *twinstar*$^{RNAi}$ egg chambers.

**(O)** Average *ATP synthase* transcription site volume in *mCherry*$^{RNAi}$ and *twinstar*$^{RNAi}$ egg chambers (n = 15).

For all plots based on imaging, each data point represents the average value for an image. Significance was assessed using a Mann-Whitney statistical test. Error bars represent standard deviation. **** P < .0001.

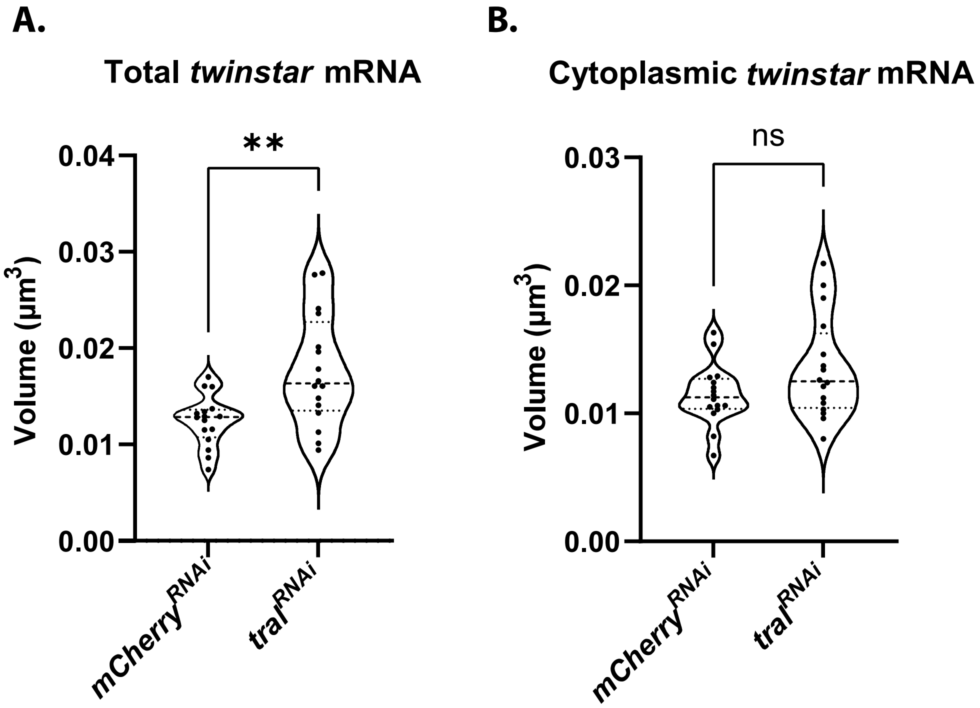


***Fig. S6 (Milano et al, 2026)***

**Figure S6: Tral contributes to the organization of *twinstar* mRNA within P-bodies.**

**(A)** Volume quantifications of *twinstar* mRNA puncta in $mCherry^{RNAi}$ and $tral^{RNAi}$ egg chambers (n = 16).

**(B)** Volume quantifications of cytoplasmic *twinstar* mRNA puncta in $mCherry^{RNAi}$ and $tral^{RNAi}$ egg chambers (n = 16).

For all plots, each data point represents the average value for all applicable *twinstar* mRNA puncta in an image. Significance was assessed using Mann-Whitney statistical tests. Error bars represent standard deviation. **** P < .0001.

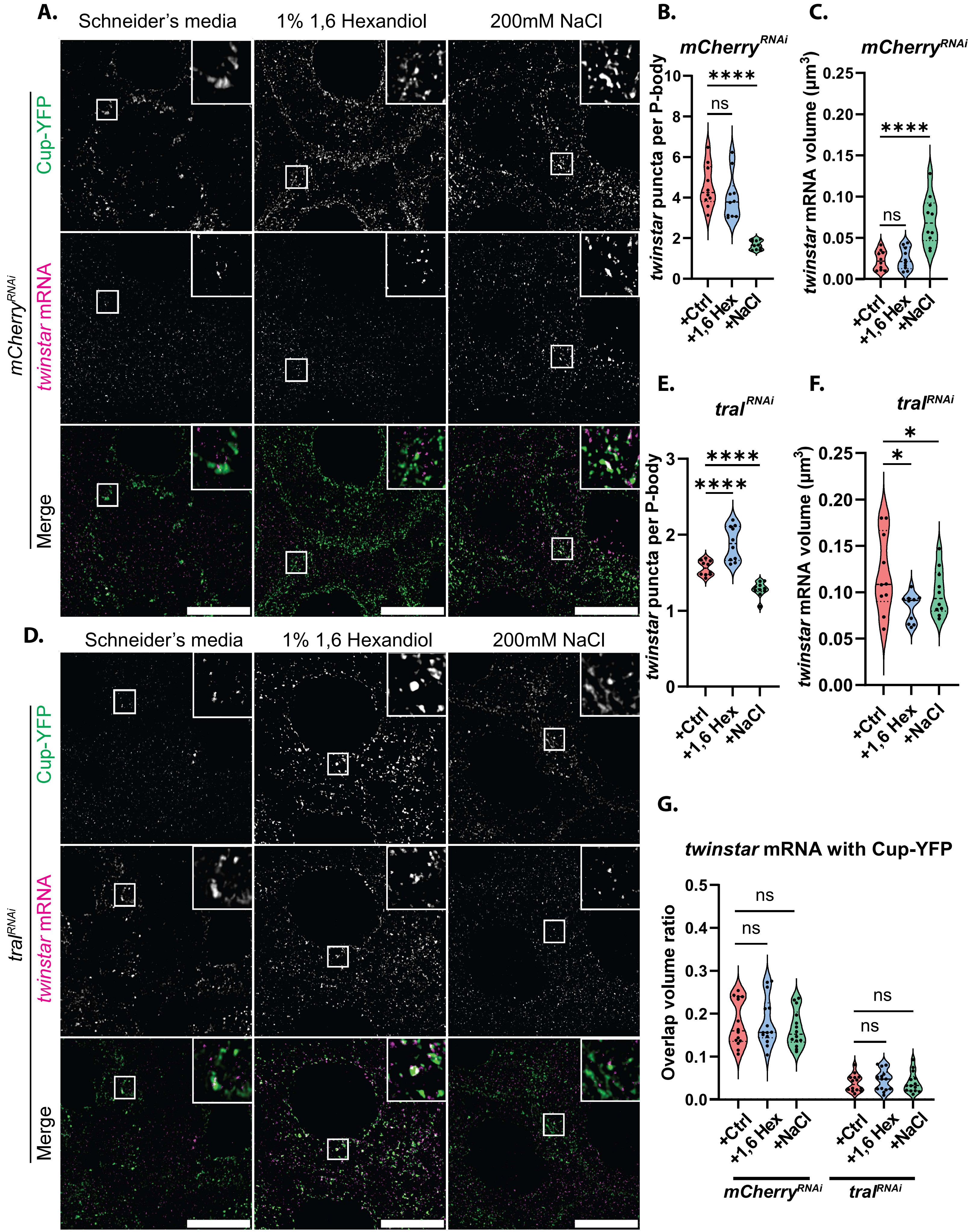


*Fig. S7 (Milano et al, 2026)*

**Figure S7: *twinstar* mRNA localization to P-bodies is dependent on association with Tral.**

**(A)** Cup-YFP visualized in $mCherry^{RNAi}$ egg chambers treated with Schneider's media (+Ctrl), 1% 1,6-hexanediol, or 200mM NaCl. Images are XY projections of 5 optical Z slices of 0.3µm. Scale bars are 20µm.

**(B)** Quantification of the average number of *twinstar* mRNA puncta per P-body in each condition in **(A)** (n = 10).

**(C)** Calculated volume of the average P-body-associated *twinstar* mRNA puncta in **(A)** (n = 10).

**(D)** Cup-YFP visualized in $tral^{RNAi}$ egg chambers treated with Schneider's media (+Ctrl), 1% 1,6-hexanediol, or 200mM NaCl. Images are XY projections of 5 optical Z slices of 0.3µm. Scale bars are 20µm.

**(E)** Quantification of the average number of *twinstar* mRNA puncta per P-body in each condition in **(D)** (n = 10).

**(F)** Calculated volume of the average P-body-associated *twinstar* mRNA puncta in **(D)** (n = 10).

**(G)** Colocalization analysis of *twinstar* mRNA puncta with Cup-YFP labeled P-bodies across conditions in $mCherry^{RNAi}$ and $tral^{RNAi}$ egg chambers.

For all plots, each data point represents the average value detected in an image. Significance was assessed using Mann-Whitney statistical tests. Error bars represent standard deviation. **** P < .0001.
